# Development and evaluation of statistical and Artificial Intelligence approaches with microbial shotgun metagenomics data as an untargeted screening tool for use in food production

**DOI:** 10.1101/2022.08.16.504221

**Authors:** Kristen L. Beck, Niina Haiminen, Akshay Agarwal, Anna Paola Carrieri, Matthew Madgwick, Jennifer Kelly, Victor Pylro, Ban Kawas, Martin Wiedmann, Erika Ganda

## Abstract

The increasing knowledge of microbial ecology in food products relating to quality and safety and the established usefulness of machine learning algorithms for anomaly detection in multiple scenarios suggests that the application of microbiome data in food production systems for anomaly detection could be a valuable approach to be used in food systems. These methods could be used to identify ingredients that deviate from their typical microbial composition, which could indicate food fraud or safety issues. The objective of this study was to assess the feasibility of using shotgun sequencing data as input into anomaly detection algorithms using fluid milk as a model system. Contrastive PCA, cluster-based methods, and explainable AI were evaluated for the detection of two anomalous sample classes using longitudinal metagenomic profiling of fluid milk compared to baseline samples collected under comparable circumstances. Traditional methods (alpha and beta diversity, clustering-based contrastive PCA, MDS, and dendrograms) failed to differentiate anomalous sample classes; however, explainable AI was able to classify anomalous vs. baseline samples and indicate microbial drivers in association with antibiotic use. We validated the potential for explainable AI to classify different milk sources using larger publicly available fluid milk 16s rDNA sequencing datasets and demonstrated that explainable AI is able to differentiate between milk storage methods, processing stage, and season. Our results indicate the application of artificial intelligence continues to hold promise in the realm of microbiome data analysis and could present further opportunities for downstream analytic automation to aid in food safety and quality.

**IMPORTANCE:** We evaluated the feasibility of using untargeted metagenomic sequencing of raw milk for detecting anomalous food ingredient content with artificial intelligence methods in a study specifically designed to test this hypothesis. We also show through analysis of publicly available fluid milk microbial data that our artificial intelligence approach is able to successfully predict milk in different stages of process-ing. The approach could potentially be applied in the food industry for safety and quality control.

## INTRODUCTION

Issues in food quality and safety can have rippling effects through the supply chain, causing substantial health and economic damage. With this, there is substantial interest in applying targeted and untargeted methods to identify ingredients or food products that show an increased risk of food fraud, food quality, and food safety issues (1, 2, 3). While targeted methods, such as the detection of toxins, pathogens, or inappropriate ingredients (e.g., horse meat in a product labeled as beef) (3), play an important role in assuring food safety and quality and preventing food fraud, they, by definition, have a set of pre-defined targets, even if (extreme) multiplexing is applied. By contrast, untargeted methods characterize all molecules that can be detected by specific method (e.g., chemical spectra, DNA sequences) to identify ingredients or products that deviate from a “baseline state” (that would be considered normal or under control) and hence would be labeled as “anomalous”. Importantly, these untargeted methods are screening methods that do not define an ingredient or product as unsafe or adulterated, rather they suggest an aberration from the normal state that should trigger follow-up actions or investigations (e.g., targeted tests, inspection of the source facility, etc.) to identify whether there are justified concerns or whether the “abnormality” detected represents natural variation that was not covered in the baseline state. While these methods can be extremely powerful to detect potential issues, they require sophisticated data analysis approaches to characterize baseline conditions and to allow for anomaly detection. In this work, bovine raw milk was selected as a model ingredient to develop improved statistical methods that can use shotgun metagenomics data as a screen to identify raw milk that shows evidence of product anomalies and deviations from baseline conditions. Milk was selected as a model as it is the sole ingredient used for the production of fluid milk— a high-volume food with considerable concern for fraud, particularly in developing countries (4). Beyond this, milk is used as an ingredient to make a variety of products and other foods, with raw milk quality having considerable impacts on finished product quality, safety, and production efficiency. Other studies have aimed to characterize the microbiome of food ingredients in production settings, for example, in high protein powders (5, 6), produce (7, 8), and fermented foods (9, 10, 11, 12). These studies are useful in demonstrating the potential that metagenomics and metatranscriptomics have in advancing food safety and quality for targeted assessments as well as for improving sensitivity for regular surveillance. Metagenomic and metatranscriptomic studies have been able to describe the microbial components of food samples with observable shifts that can be related back to key attributes of metadata, e.g., ingredient contamination (5). However, it should be noted that when studies rely on amplicon sequencing (often due to cost and resource limitations), there can be important reductions in sensitivity and taxonomic resolution (13).

The food supply chain is highly complex, with a multitude of touch points (e.g. farmers, suppliers, transportation, storage, etc.) existing prior to reaching a finished product, where issues occurring at each step have the potential to cause quality and safety issues. Therefore, while these early studies have set a foundation for the use of the microbiome in food production, the expansion of these analyses and applications into additional food ingredients and for different supply chain challenges will only continue to refine and increase the robustness of analysis similar to how much work has been done to build microbiome standards across human-associated (14, 15, 16, 17) and environmental niches (18).

Our objective was to expand on the growing evidence that the microbiome can be used as a relevant indicator through an application in fluid milk with the hypothesis that it could be used to identify (i) raw milk that represented a farm different from a given, predefined source farm (simulating introduction of an unknown or unapproved supplier into an ingredient stream - “outside farm”, abbreviated as OF) or (ii) raw milk that contains some milk from mastitis-affected cows treated with antibiotics (representing a regulatory violation - “antibiotic treated” abbreviated as ABX). Importantly, these scenarios were simulated by using commercially produced milk without applying any additional *a priori* targeted testing to assure that these “anomalous samples” show easily identifiable differences from the baseline samples. This approach was used to provide a real-world and realistic dataset for the proof-of-concept study described here.

Milk was used as a model system for examining the application of metagenomic sequencing of microbial communities for food safety and quality, building on our earlier work evaluating associated DNA extraction and host depletion methods (19). Raw milk microbiomes have been found to be diverse (20, 21, 22, 23) and potentially have an influence on the quality of downstream processed dairy products (24, 25). These and other published works support utilizing milk microbiomes as a potential source of information for quality assurance and risk assessment in the food industry.

We evaluated different anomaly detection methods beginning with the classical microbiology ecological metrics of alpha and beta diversity, differential abundance, clustering, as well as ordination through contrastive PCA and MDS. However, these classical methods were limited in their ability to differentiate sample classes. In turn, a growing number of studies have demonstrated the benefit of leveraging machine learning to differentiate sample classes in microbiome studies. These include predicting the risk of type 2 diabetes (26), diarrhea associated with cancer treatment (27), and liver disease (28) from the gut microbiome. Additionally, when sampling the microbiome from human skin, explainable AI was able to identify microbial drivers associated with skin hydration, age, and pre/post-menopausal status from the skin microbiome (29).

For this work, we collected 58 bulk tank milk samples in a block-randomized time-constrained design to assess the ability of the microbiome to indicate deviations from a baseline (BL) community related to anomalies (outside farm and antibiotic use) that could be present in the food supply chain and be related to food quality issues. A set of 33 consensus microbes were found to be stable elements in baseline shotgun metagenomics samples with *Pseudomonas, Serratia, Cutibacterium*, and *Staphylococcus* to be the most abundant. Traditional methods of ordination (cPCA and MDS), as well as alpha and beta diversity, were limited in their ability to fully separate sample classes and microbial differences associated with anomalies. However, explainable AI was able to differentiate sample classes while also identifying three key microbial drivers that separated sample classes with significance even in this dataset, which would be considered small in the realm of machine learning techniques studies. Given that whole genome shotgun sequencing is still prohibitively costly for wide application, we next investigated if other datatypes could be used. We applied explainable AI to 16S rRNA data from two publicly available milk microbiome datasets to confirm that this ap-proach could distinguish between milk from different categories. We demonstrate that our explainable AI approach is able to successfully predict the processing stage and the transport stage a milk sample comes from. This study provides advances in the application of machine learning that can be expanded across the food industry.

## RESULTS

### Collection and shotgun metagenome sequencing of baseline and anomalous raw milk samples

For whole metagenome shotgun sequencing, in total 65 samples were collected: 33 baseline (BL), 13 outside farm (OF), 12 antibiotic treated (ABX), 6 negative DNA extraction controls, and 1 sequencing blank. Anomalies were chosen to represent potential sources of concern in a dairy processing plant (milk from an unknown source or milk contaminated with antibiotic residues). The sampling scheme is shown in Supplemental Figure S1.

Sampling dates were block-randomized to ensure even distribution across the sampling period for each anomaly type. A short time frame of five weeks was chosen to control for seasonality.

Metadata including milk components (e.g., lactose, fat, milk urea nitrogen), somatic cell count, and standard plate count were also collected and provided as Supplemental File S1. No strong correlations were observed between the metadata features and individual microbe reads per million (RPM) values or the summed microbe RPM values per sample (|Spearman corr.| < 0.7).

Shotgun whole metagenome sequencing (Methods Section 3.2) with Illumina NovaSeq 6000 at 2 × 150bp resulted in 39.6-79.2 million read pairs per raw milk sample. Figure 1 details the overall bioinformatics pipeline that was applied to this data.

**FIG 1.**
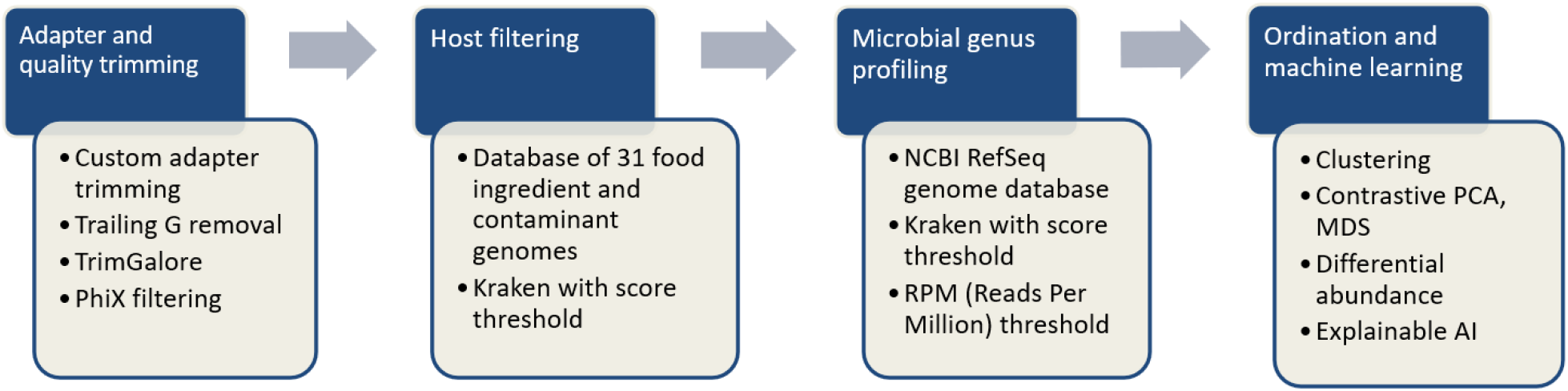
Overall analysis pipeline.

During quality control of the reads with FastQC (30) and manual inspection, full-length junction adapters and trailing G’s were observed, indicating that the DNA was likely fragmented prior to sequencing. These artifacts were removed (Methods Section 3.3), and resulting reads were trimmed for low quality bases using TrimGalore (31). A median of 59.3M read pairs per sample were retained as high quality with a median of 0.5% read pairs needing removal due to quality issues (see Supplemental Figure S2 and Supplemental Table S2).

To remove bovine and potential contaminant sequence content, we employed matrix filtering as reported in a recent publication on food microbiome sequencing (5). Kraken (32) was utilized with a custom-built reference database of 31 common food ingredients and contaminant genomes (5) including *Bos taurus* (assembly Btau_5.0.1) and *Homo sapiens* (assembly GRCh38.p10). A large fraction, 91.3–98.7%, of reads were discarded from subsequent microbial analysis as matrix-classified (see Supplemental Figure S2(a) and Supplemental Table S2). Negative sequencing controls were utilized to quantify the presence of typical laboratory contaminants that can be expected in shotgun sequencing as it has been previously reported (33), particularly in studies using low biomass samples (34, 35, 36). With this, the background microbial contamination was identified bioinformatically using the decontam (37) R package on the genus-annotated reads. The analysis identified 14 genera which were removed from subsequent analysis: *Histophilus, Rahnella, Raoultella, T4virus, Pragia, C2virus, Methylophilus, Oceanobacillus, Streptosporangium, Fluviicola, Oenococcus, Alkalilimnicola, Geminocystis*, and *Brevibacillus*.

### Microbiome characterization of raw bulk tank milk

We utilized Kraken (32) with the NCBI RefSeq (38) whole genome collection to annotate the high quality non-matrix read pairs and summarized the classified reads at genus level (Supplemental Table S3). Each genus observed was defined based on a minimum abundance of 0.1 Reads per Million (RPM) as indicated in (5), resulting in 572 observed genera.

Alpha diversity was determined using the Shannon index calculated based on the genus level table. Four baseline and two outside farm samples (BL-04, BL-08, BL-13, BL-16, OF-09, and OF-13) were observed to have very low diversity (Shannon index <0.4, Figure 2a), with a high relative proportion of reads assigned to specific microbes (Supplemental Figure S2(b)), indicating a potential ‘bloom’ of specific milk microbes in those samples, which drove diversity indexes to extremely low levels. This was further confirmed by the observation that these samples were dominated by a single organism where 94–95% of annotated reads were classified as either *Pseudomonas* or *Staphylococcus* (in the case of OF-13). While *Pseudomonas* and *Staphylococcus* have been previously detected in milk, the observation of a single organism accounting for the vast majority of the microbial profile of a sample would most likely be due to a random sample-specific event, and not milk microbiome signatures associated with a given farm. While these six samples were therefore removed from further analyses, once larger datasets are available for analyses, samples with these types of sporadic ‘blooms’ would not need to be removed, as they could be identified algorithmically as outliers.

**FIG 2.**
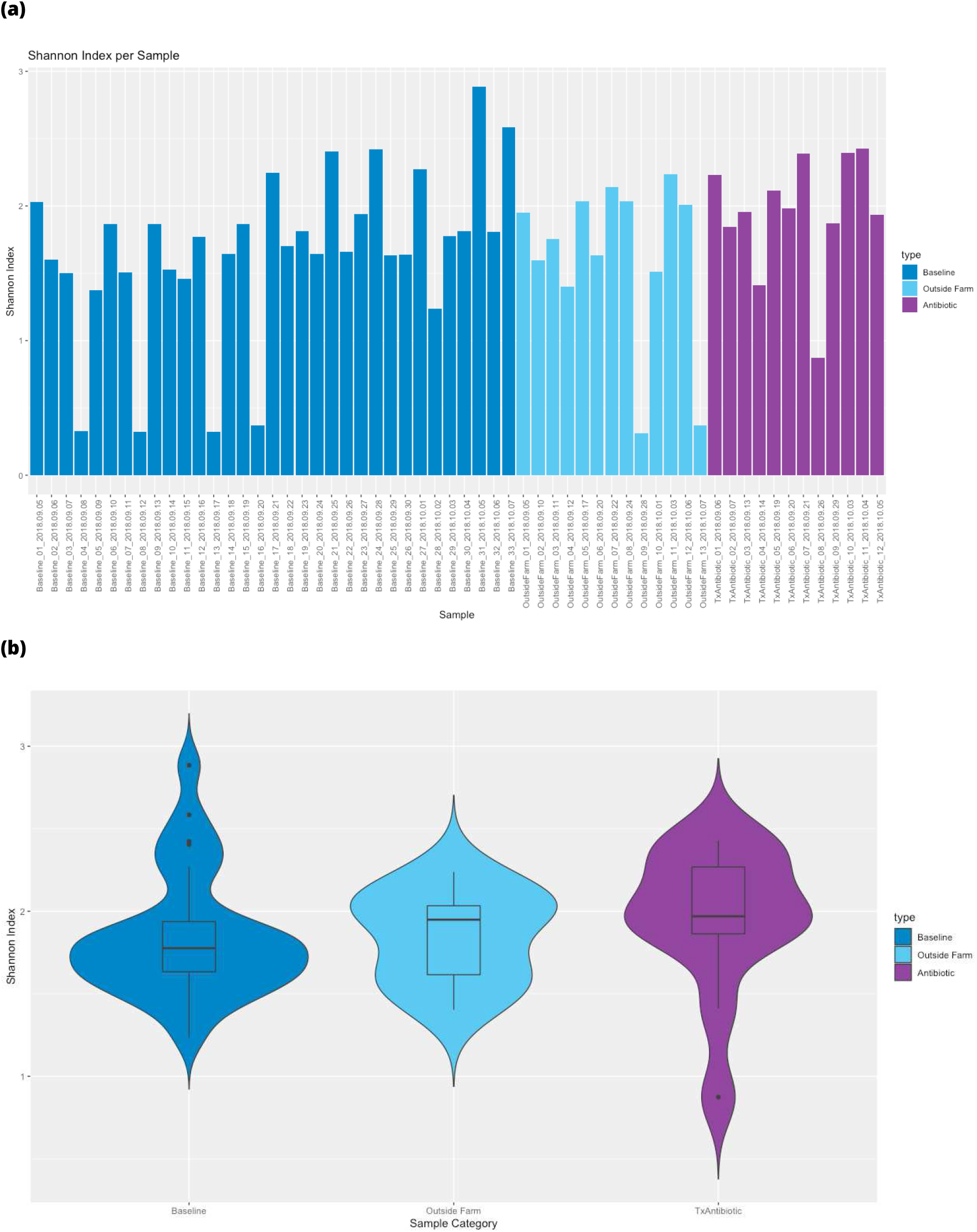
Alpha diversity (Shannon index) of all raw milk samples where presence is indicated as genus RPM > 0.1. **(a)** Shannon index is shown for all 58 samples and **(b)** summarized by sample class with the six low-diversity outlier samples removed here and in all subsequent figures (“Baseline_04”, “Baseline_08”, “Baseline_13”, “Baseline_16”, “OutsideFarm_09”, “OutsideFarm_13”).

The remaining samples had an average of 27K microbial classified reads at genus taxonomic rank or more specific. The alpha diversity within each sample class was relatively consistent (Figure 2). The average Shannon index per sample class was 1.84–1.95 with an average number of genera observed per sample class to be 143–162 with no major differences due to sampling date. Between sample classes, the alpha diversity shows a similar distribution (Figure 2b) with a Wilcoxon rank sum test for BL vs OF *p* = 0.7654 and BL vs ABX *p* = 0.1194.

Classified reads per million quality-controlled sequenced reads (RPM) were computed for each genus, and a threshold of 0.1 RPM was applied to define *supported* genera, as described in Beck et al. (5). The supported genera RPM values are provided as Supplemental Table S4. Figure 3 highlights the most abundant microbes observed in the milk microbiomes. Genera with RPM greater than 5% of the supported genera RPM total in at least one sample are shown (with remaining genera summed as “Other”). In total, there were 12 such genera, and combined they account for 73.5–97.3% of the total supported genera RPM sum per sample. Only *Pseudomonas, Serratia, Cutibacterium*, and *Staphylococcus* were observed to account for more than 5% of the total supported genera RPM in every sample.

**FIG 3.**
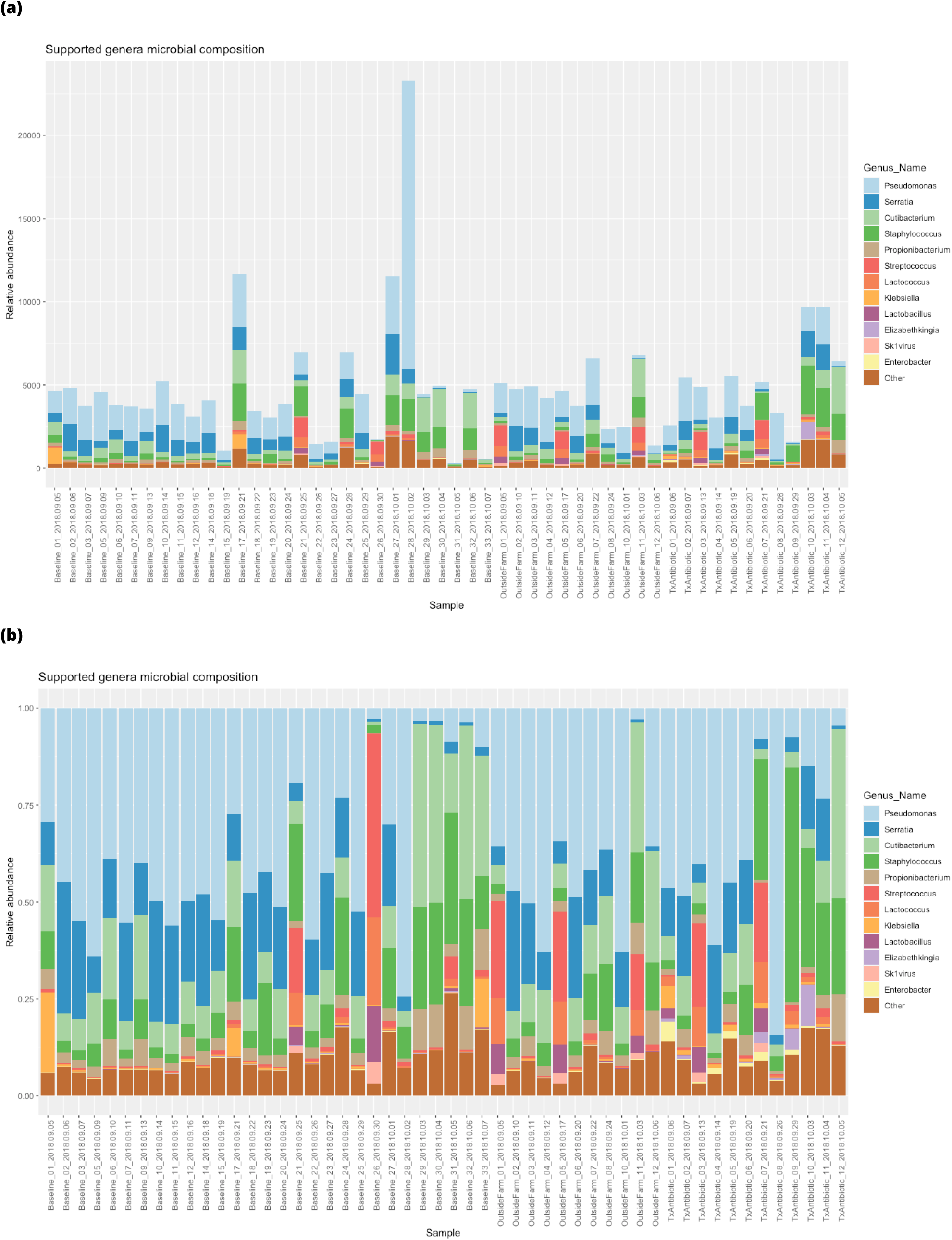
Microbial community membership is shown per sample for genera with RPM abundance greater than 5% in any sample in (**a**) standard stacked barplot and (**b**) with values scaled to 100% by sample. Genera observed in less than 5% of the summed per sample abundance are aggregated into the “Other” class.

Additionally, a date-localized increase in abundance of *Cutibacterium* from 03-Oct. to 07-Oct. was observed in baseline, outside farm, and antibiotic treated samples. The anomalous samples were from sub-sampled dates based on our block random-ized sampling selection (in comparison to continuous date sampling for the baseline). Additionally, there were also a select number of microbes observed in only a few samples within one class. For example, *Klebsiella* was only observed in BL-01, BL-17, and BL-33 as high abundance, and *Enterobacter* was uniquely observed in antibiotic treated samples (ABX-01) as was *Elizabethkingia* (ABX-10, ABX-11).

### Contrasting baseline and anomalous community profiles using traditional methods

To contrast sample classes and better understand microbial drivers associated with anomalous sample classes, we began by exploring traditional microbial ecology and ordination methods.

Beta diversity was calculated by abiding by principles of compositional data analysis and computing an Aitchison distance as described in Beck et al. (5) and in Methods Section 3.5. This distance was used to cluster samples as shown in Figure 4a. The samples clustered into three main clades with intermixed sample classes and sampling dates. The first clade (furthest left, OF-11, BL-29, BL-32, BL-30, ABX-12) was driven by the presence of *Propionibacterium*, but not with the co-occurrence with *Klebsiella* as in BL-33. The remaining two larger clades intermixed all sample types and dates without notable microbial differences defining their structure and separations.

**FIG 4.**
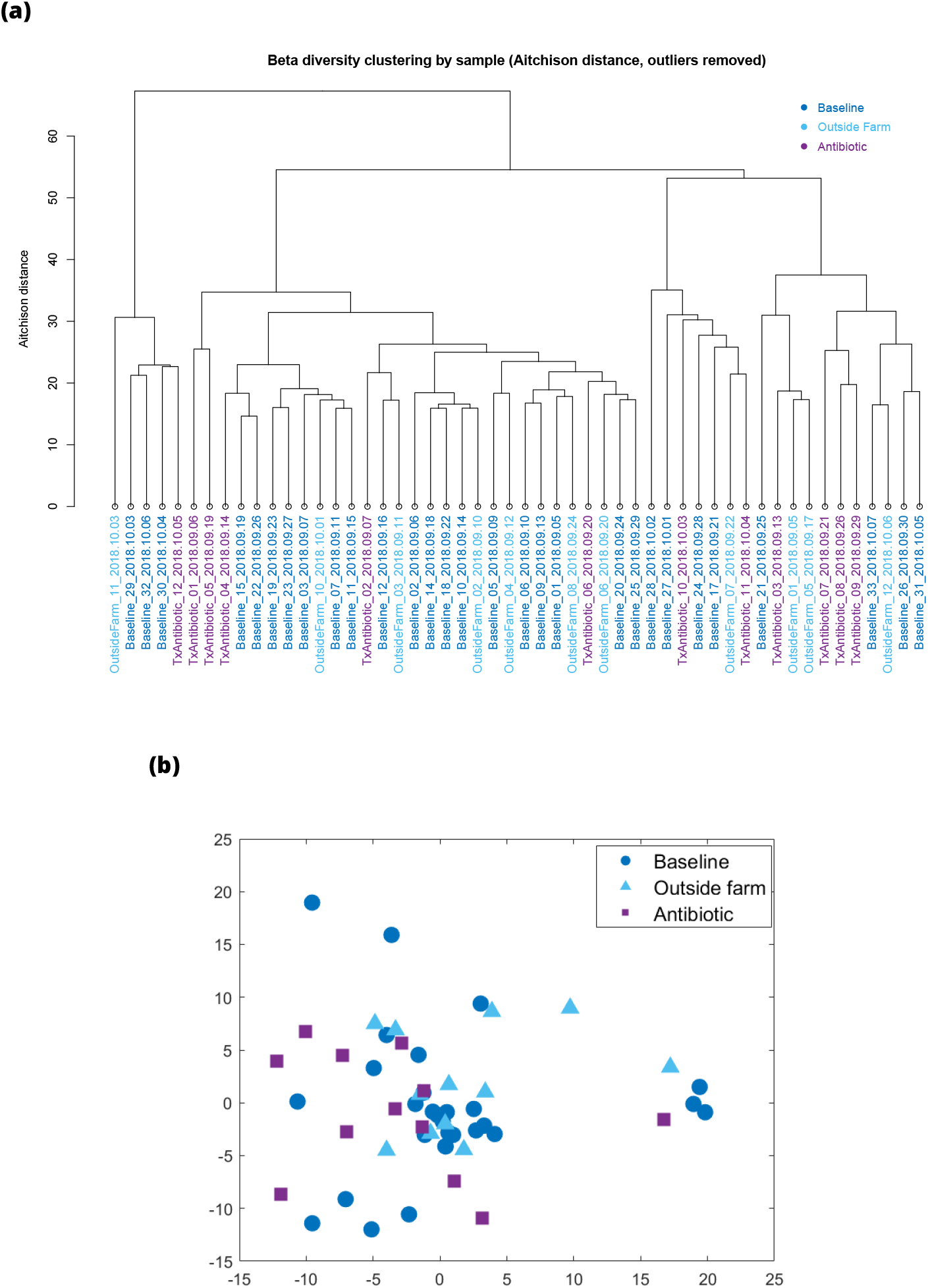
Beta diversity using Aitchison distance of raw milk samples with six outlier samples removed is shown here in **(a)** hierarchical clustering and **(b)** multidimensional scaling.

Contrastive PCA (cPCA) (39) was applied to the microbial community relative abundance data. However, it was not able to successfully separate the outside farm, antibiotic, and baseline samples when baseline samples were used as the background dataset (Supplemental Figure S3). The difference between PCA and cPCA is that cPCA aims to identify enriched patterns in a target dataset (foreground dataset) by contrasting it with another dataset (background dataset). This is an unsupervised technique that uses a hyperparameter named alpha to adjust the trade off between high target variance and low background variance. At alpha = 0, cPCA collapses to PCA on the target dataset. At alpha = inf, it puts an infinite penalty on any direction which is not in the null space of the background dataset.

In contrast to cPCA which operates on the microbial RPM count table, multidimensional scaling ingests a pairwise distance matrix and aims to project the samples onto a lower dimensional space while retaining their distances. Multidimensional scaling (MDS) based on the Aitchison distance indicated some separation between the anomalous sample types ABX and OF in the two-dimensional projection (Figure 4b), although the baseline samples appear intermixed with the anomalous samples. The three classes are significantly separated according to PERMANOVA (*p* = 0.0064).

### Differentially abundant genera

Two-sample Kolmogorov-Smirnov tests were performed independently for each genus to determine statistically significant differentially abundant features. After Bonferroni correction for multiple testing, the adjusted *p*-values were significant (*p* < 0.01) for *Coxiella* for BL vs. OF and *Enterobacter, Morganella* for BL vs. ABX. Their RPM distributions are visualized in Figure 5. *Coxiella* was observed to be increased in outside farm samples and *Enterobacter* and *Morganella* in antibiotic treated samples. Most notably, the median RPM value of *Enterobacter* in ABX samples was nearly 12 times that of the baseline samples (28.0 vs. 2.4 RPM).

**FIG 5.**
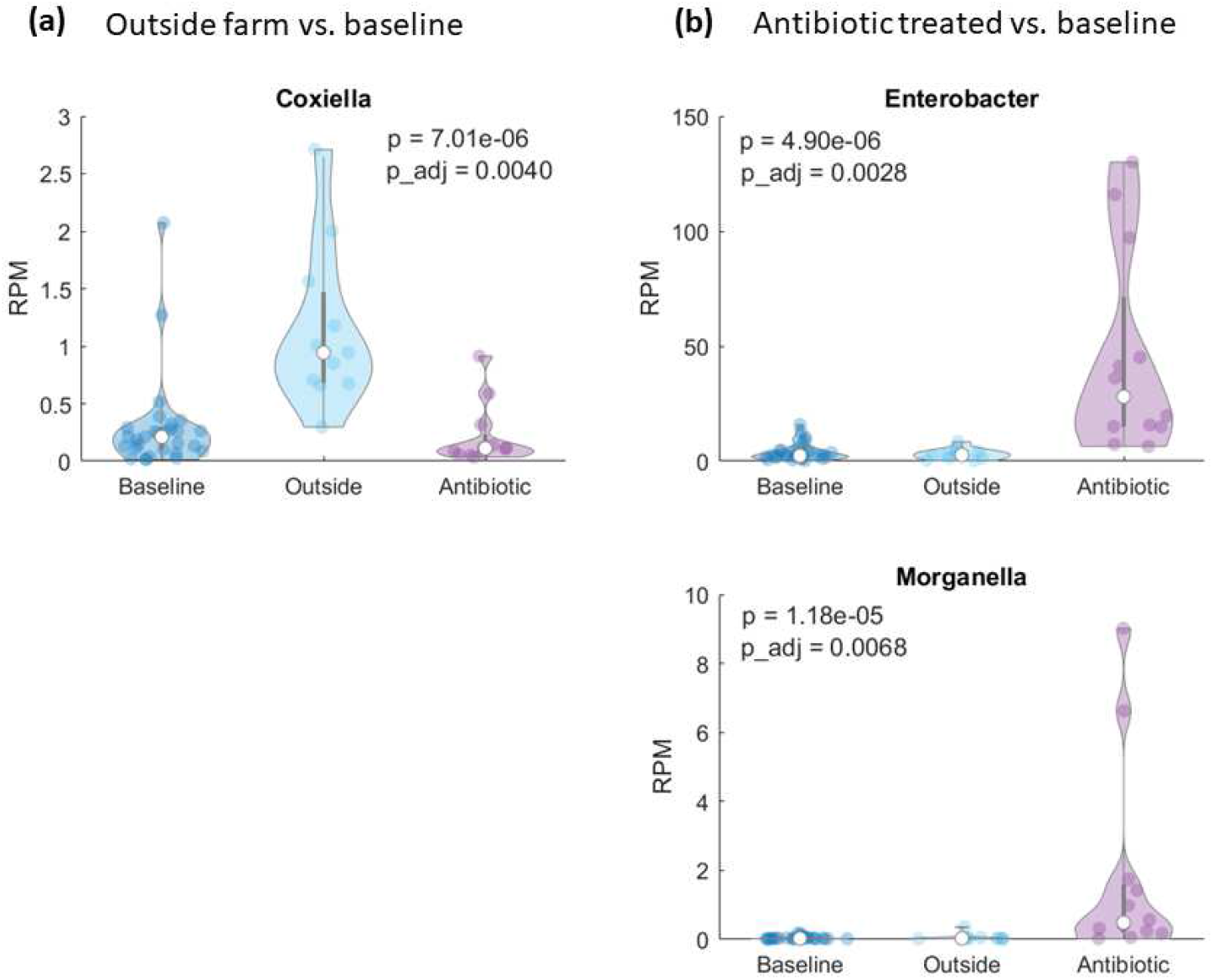
Distribution of RPM values for each group, for the differentially abundant genera. **(a)** Outside farm vs. baseline samples. **(b)** Antibiotic treated vs. baseline samples. Unadjusted and adjusted p-values from the two-sample KS-test are shown within each plot. White circles indicate median values.

### Sample class prediction with explainable AI

We employed an explainable AI workflow (‘AutoXAI4Omics’ (https://github.com/IBM/AutoXAI4Omics)), as described in a recent study on the skin microbiome (29), to perform two separate classification tasks: BL vs. OF and BL vs. ABX. The genus-level RPM data after removing low-diversity outlier samples and contaminant genera was used as input (52 samples, 572 features). For each classification task, the samples were split into training (70%) and test (30%) sets uniformly at random, while maintaining the class size distribution in each set. Five randomized iterations of the train/test split were performed to obtain robust results. To evaluate and compare the predictive performance of our machine learning model, we used a state-of-the-art measure of accuracy, the F1-score. The F1-score is the harmonic mean between precision and recall. It is a metric of accuracy that takes into account the imbalance of the classes when the average parameter is set to “weighted”.

The AutoXAI4Omics workflow included training and tuning of several machine learning algorithms (XGboost, Random Forest (RF), Support Vector Machines, Adaboost, K-Nearest Neighbors (KNN), LightGBM, Decision Trees, Extra Trees, Gradient Boosting, Stochastic Gradient Descent) followed by selection of the best model based on the predictive performance represented by the F1-score. The three best machine learning models were tree-based models (XGBoost, Random Forest, LightGBM). XGBoost however, a gradient-boosted decision tree ensemble method, consistently reported a higher F1-score on the test data and during cross validation. XGBoost has also performed well in recent comparative studies on microbiome data (40, 29, 41).

Generating explanations for the ML models’ predictions is an important active field of research. Understanding and explaining the mechanisms underlying the predictions can help validate the predictive models and reveal powerful and novel insights about the connections between the samples and phenotype under investigation. Therefore, for each classification task, we used AutoXAI4Omics to generate explanations of the predictions for XGBoost using an explainable AI algorithm called SHapley Additive exPlanations (SHAP) (42). The SHAP algorithm assigns a SHAP value to each genus (i.e., feature) that represents the impact, negative or positive, that the genus has in predicting a class for a given sample. The genera are then ranked based on their average absolute SHAP impact value across all the samples in the training dataset to obtain a ranked list of impactful genera (see Methods Section 3.7 for more details).

Given the number of samples per class, the random variability in the splitting of samples into training and test sets has an effect on the predictive performance and the ranked list of the most impactful genera (i.e., when running multiple iterations while changing the global random seed). As such, the predictive performance and the stability (43) of the top impactful genera from SHAP were examined across five randomized iterations. We observed overall high variation in the order of the impactful genera across the five randomized iterations with different random seeds. *Stable predictive features* were defined as those being among the top three most impactful features in at least two out of five randomized iterations.

Despite the observed variation, three stable impactful features were identified: *Coxiella* for the BL vs. OF comparison as well as *Enterobacter* and *Morganella* for the BL vs. ABX comparison. These stable features that are impactful for the prediction are the same as the differentially abundant genera identified above with the KS-test.

In addition to confirming the predictive impact of the three statistically significant genera, one advantage of the explainable AI algorithm, SHAP, over other feature importance methods, is that it also explains how each of these impactful features is contributing, positively or negatively, to the prediction of a particular class (e.g., BL) for each sample or across a set of samples.

In Figure 6(a) for one of the five runs, *Coxiella* is shown as positively contributing to the prediction of the class OF for those samples that have a higher abundance of *Coxiella*. Similarly, in Figure 6(b) *Enterobacter* and *Morganella* are driving the prediction of ABX for those samples with higher abundances of the genera. The SHAP plots for the remaining four iterations are shown in Supplemental Figure S4.

**FIG 6.**
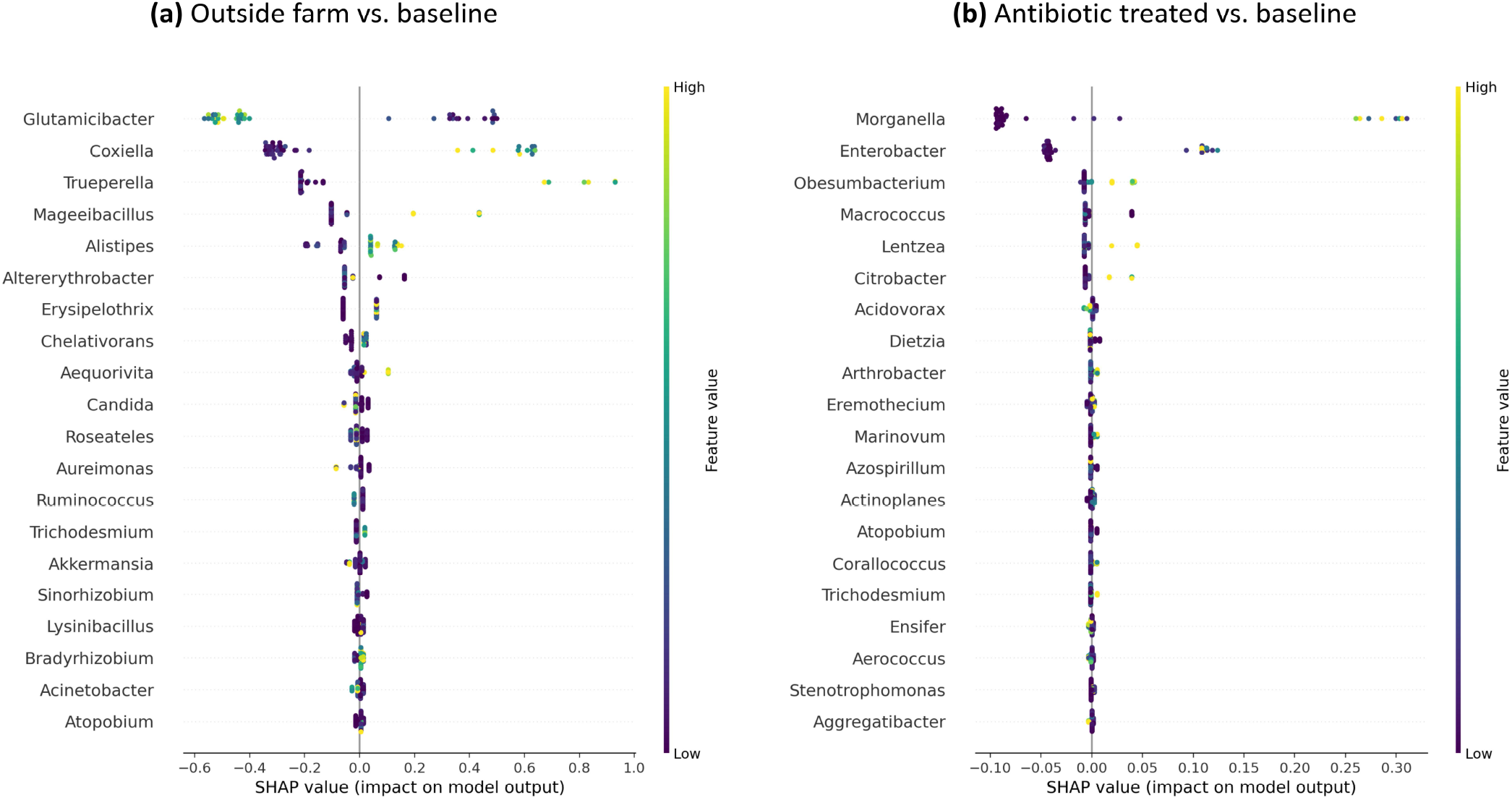
Explainable AI results, SHAP summary dot plots in one exemplar iteration of each anomaly type comparison against baseline (see Supplemental Figure S4 for the other four iterations). SHAP values indicate the importance of that feature on the prediction of the sample class (see further explanation in Section 3.7). **(a)** The most impactful features predicting outside farm anomaly class. **(b)** The most impactful features predicting antibiotic treated anomaly class.

For the BL vs. OF prediction, in 4/5 runs *Coxiella* was the most impactful and in 1/5 runs it was the second most impactful. The F1-score of BL vs. OF prediction ranged from 0.833 to 0.917 (mean 0.87). For the BL vs. ABX prediction, in 4/5 runs *Morganella* was the most impactful. In 1/5 runs *Enterobacter* was the most impactful and in 2/5 runs the second most impactful. The F1-score of BL vs. OF prediction ranged from 0.692 to 0.923 (mean 0.83).

### Validation of explainable AI approach using alternate datatype

To validate the ability of using an explainable AI approach to detect anomalous milk samples based on microbiome composition, next we applied this technique to an alternative datatype. We selected 16S rRNA data, an affordable and accessible proxy for whole genome shotgun sequencing metagenomics. We selected two publicly available fluid-milk microbiome datasets (ERP015209, ERP114733), containing 1,507 and 626 samples with 16S rRNA data, respectively (23, 20). We then used the AutoXAI4Omics tool to investigate whether 16s rRNA data could also distinguish milk from a range of categories, including season, transport stage, silo ID, and processing stage. For processing stage comparisons, classes included raw milk stored in silos “Raw Milk” class, milk stream entering the pasteurizer “HTST feed” class, and post-pasteurization milk “HTST Milk” class. For transport stage comparisons, classes included milk that was collected with a stainless steel dipper from the inlet at the top of individual tanker trucks “Tanker” class, raw milk sampled from five large-volume-capacity silos “Raw Milk” class, and “Blended Silo” class.

Of the six ML models tuned, trained, and cross-validated by AutoXAI4Omics (RF, KNN, AutoKeras, LightGBM, Autosklearn and XGBoost), RF performed the best, predicting processing stage with 0.734 F1 score (Table 1). SHAP analysis revealed that the most impactful genera influencing prediction of raw milk samples included lower abundances of the genus *Bacillus* and the thermophilic genus *Thermus*, and higher abundances of genera *Pseudomonas* and *Acinetobacter* (Figure 7). After the pasteurization process (high-temperature short-time processing (HTST)), the genera listed above appeared to have an opposing influence upon the model’s prediction. We observed that in pasteurized samples, higher abundances of *Bacillus* and *Thermus*, and lower abundances of *Pseudomonas* and *Acinetobacter* influenced the prediction of this class (Figure 7).

**TABLE 1.** Explainable AI performance on alternative datasets. The average F1 score and standard deviation of the best performing model after 5-fold cross-validation on each of the alternative datasets are indicated with the number of training and test samples.

| Target Variable | Best Model | Train:Test | F1-Score Train | F1-Score Test | Study ID |
| --- | --- | --- | --- | --- | --- |
| Processing Stage | Random Forest | 311:78 | 0.997 ± 0.028 | 0.734 ± 0.040 | ERP114733 (23) |
| Transport Stage | Random Forest | 303:76 | 0.998 ± 0.021 | 0.885 ± 0.013 | ERP015209 (20) |
| Season | XGBoost | 986:298 | 0.998 ± 0.002 | 0.849 ± 0.023 | ERP015209 (20) |

**FIG 7.**
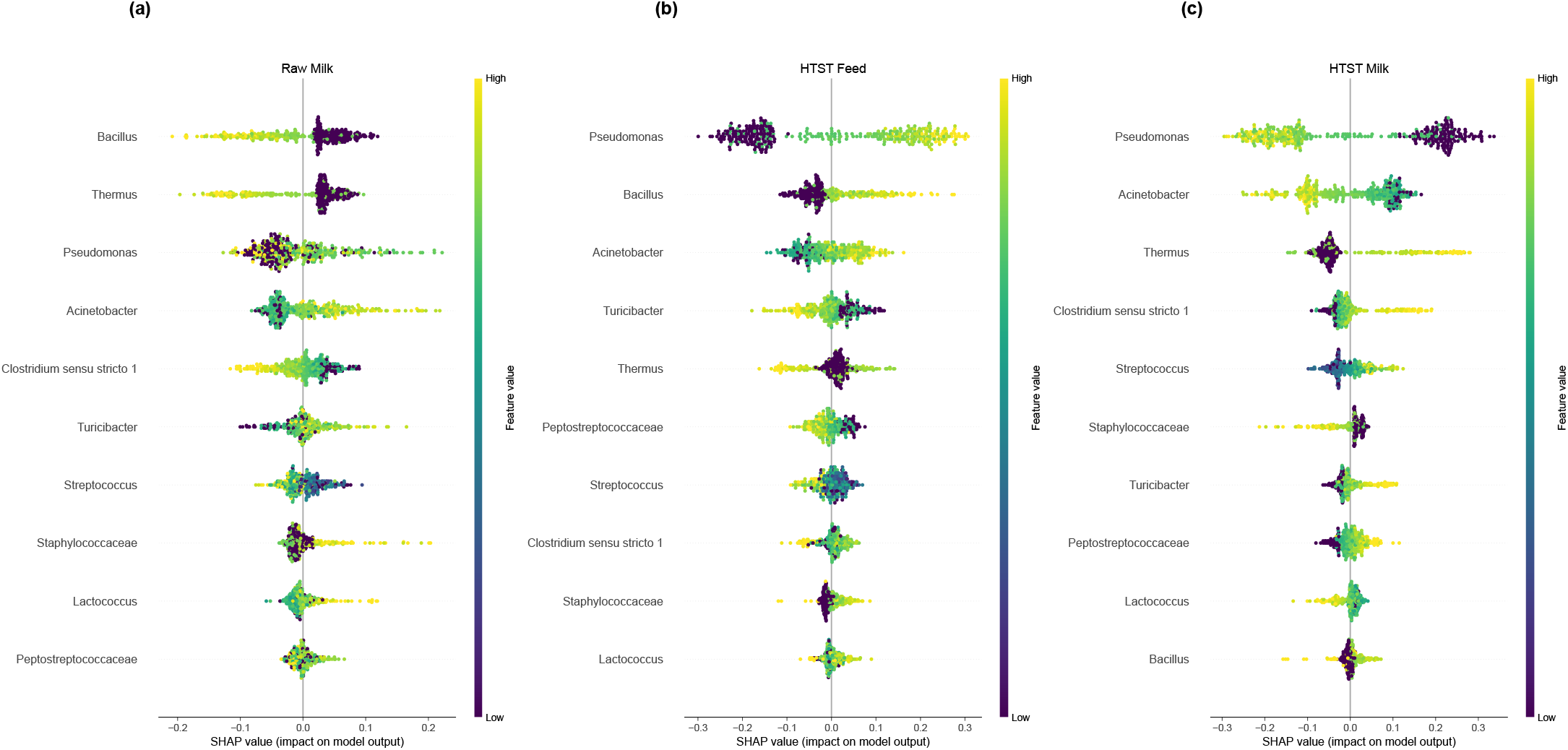
Explainable AI results on validation with alternative datatype, evaluating the ability to predict the processing stage (n samples train:test were 311:78). SHAP summary dot plots in one exemplar iteration of each anomaly type comparison. SHAP values indicate the importance of that feature on the prediction of the sample class (see further explanation in Section 3.7). **(a)** The most impactful features predicting raw milk class. **(b)** The most impactful features predicting pre-pasteurization class. **(c)** The most impactful features predicting post-pasteurization class.

Another category for which AutoXAI4Omics produced highly accurate models was milk storage stage, which represented different locations within the milk transport pipeline, raw milk, tanker milk, or silo milk. The best performing model predicting milk transport stage was Random Forest, with an average F1-score of 0.885 across the three classes (Table 1). For tanker milk and silo milk, *Bacillus, Mycoplasma*, and *Lactococcus* were highly influential in the model’s prediction, but they showed opposing influences for each of the two classes. Lower abundances of *Bacillus, Mycoplasma*, and *Lactococcus* were associated with tanker milk prediction, whilst higher abundances of these two genera influenced silo milk prediction (Figure 8).

**FIG 8.**
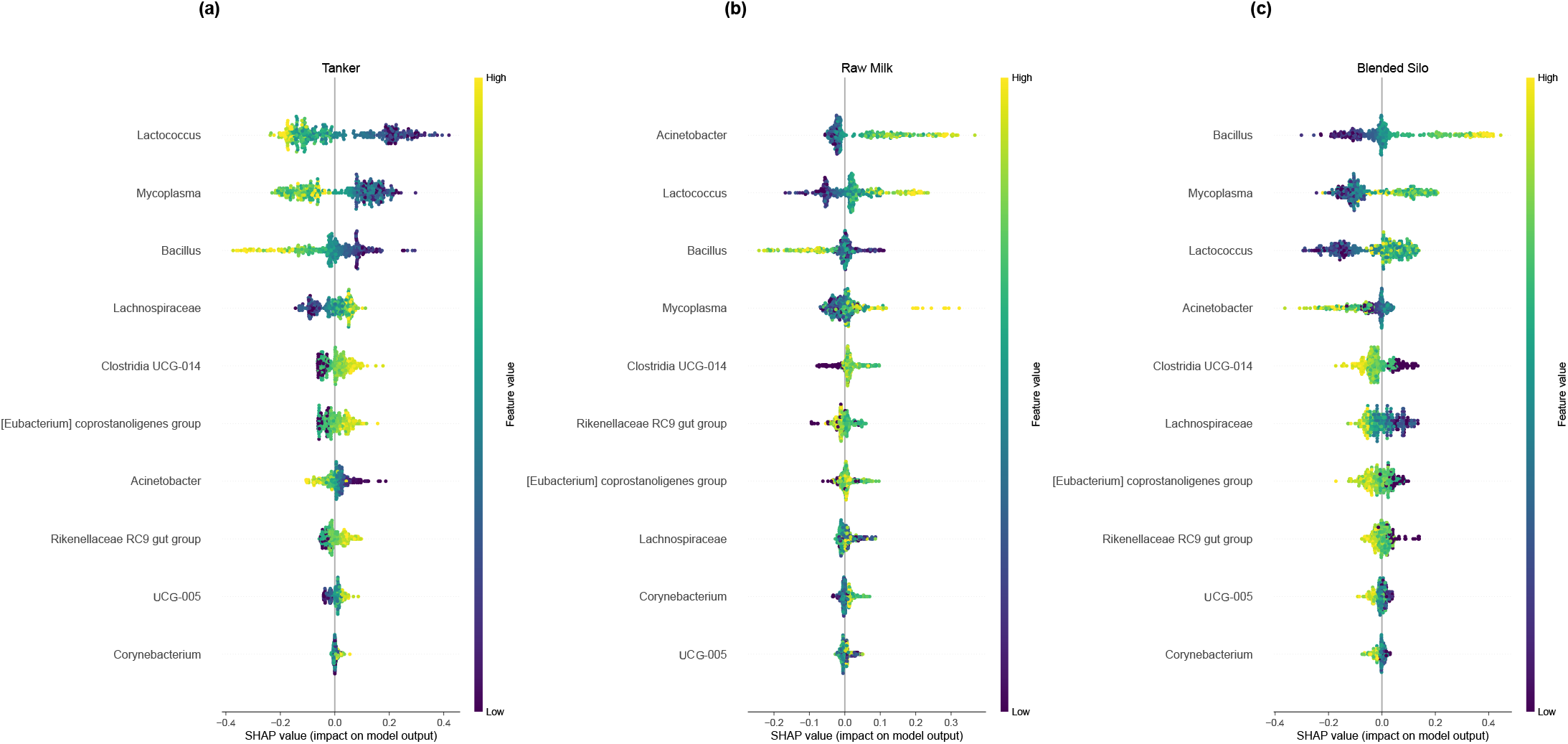
Explainable AI results on validation with alternative datatype, evaluating the ability to predict milk storage stage during transport (n samples train:test were 303:76). SHAP summary dot plots in one exemplar iteration of each anomaly type comparison. SHAP values indicate the importance of that feature on the prediction of the sample class (see further explanation in Section 3.7). **(a)** The most impactful features predicting tanker milk class. **(b)** The most impactful features predicting raw milk class. **(c)** The most impactful features predicting blended silo class.

AutoXAI4Omics was also able to successfully predict the season in which a milk sample was collected, using categories Fall, late Summer, Summer and Spring (Table 1). XGBoost predicted season with the highest accuracy of all models, with an average F1-score across the four classes of 0.849 (Table 1) *Mycoplasma* most strongly influenced prediction of both late Summer and Fall classes; however, a higher abundance increased the likelihood of a late-Summer prediction, whilst a lower abundance influenced classification as a Fall sample (Figure 9). *Mycoplasma* was also the second most influential genera in the prediction of a Spring sample, suggesting this genus is heavily influenced by season.

**FIG 9.**
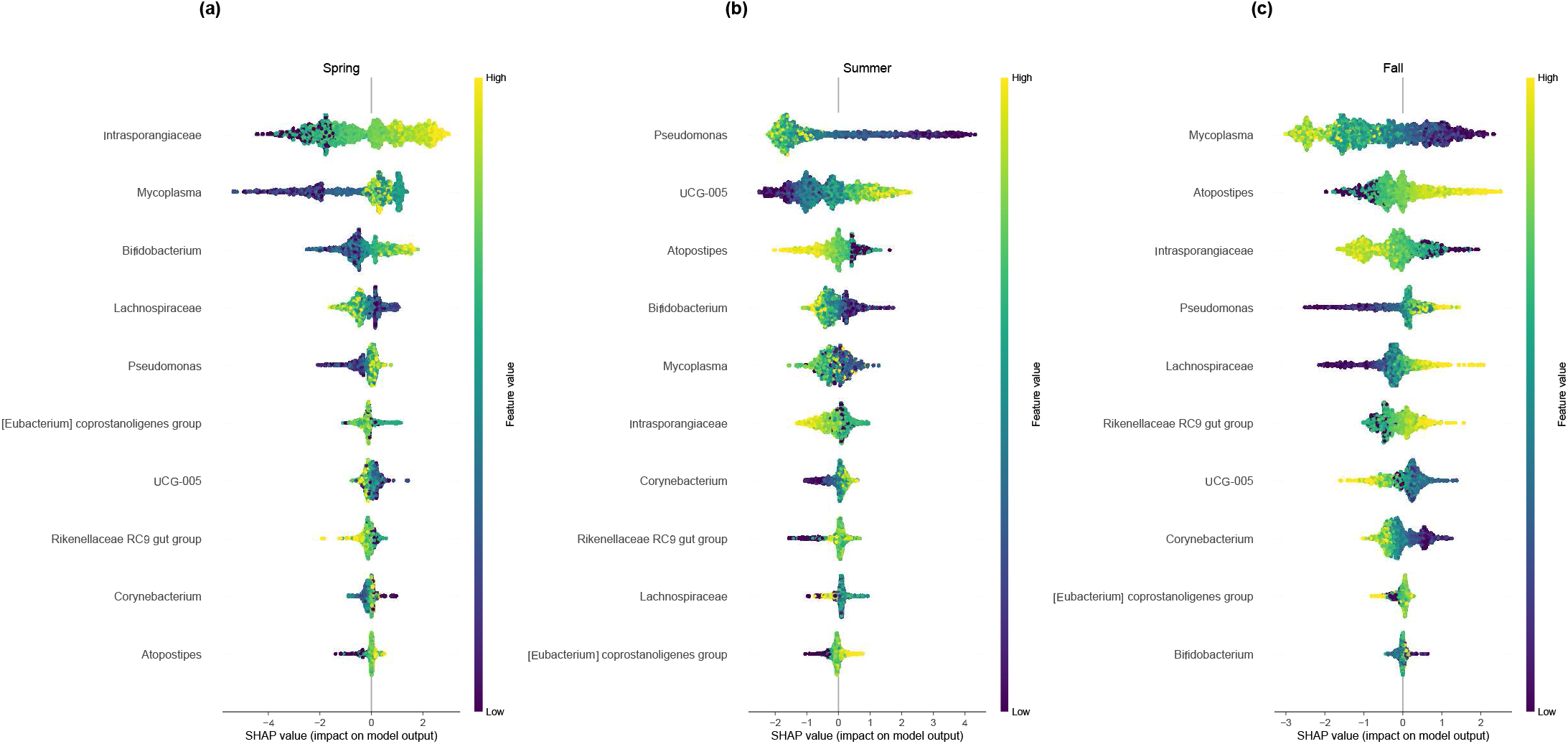
Explainable AI results on validation with alternative datatype, evaluating the ability to predict milk collected in different seasons (n samples train:test were 986:298). SHAP summary dot plots in one exemplar iteration of each anomaly type comparison. SHAP values indicate the importance of that feature on the prediction of the sample class (see further explanation in Section 3.7). **(a)** The most impactful features predicting spring class. **(b)** The most impactful features predicting summer class **(c)** The most impactful features predicting fall class.

Milk samples collected from different silos could not be accurately predicted using our explainable AI approach (F1-score 0.25), indicating limited differences in 16S rRNA microbial composition between milk stored in different silos. This finding is commensurate with findings by Kable et al., (20) observing no clear difference using hierarchical clustering of beta diversity of samples.

## DISCUSSION

We have applied various traditional and artificial intelligence methods to test the hypothesis that the microbial community of raw milk, as characterized by shotgun metagenomics, could be used for anomaly detection based on the intuition that the microbiome would differ in relation to two anomalous states: (i) treatment with antibiotics and (ii) the presence of milk from a differing farm. The rationale for testing such anomalous states was to attempt to detect anomalies that would be meaningful to the industry (e.g., regulatory violation or unknown ingredient source) (44, 45). The microbiome has been shown to be a highly dynamic ecosystem where microbial community membership and relative abundances can shift in response to a variety of perturbations (46). This has been demonstrated in health (47), the environment (48), and more recently in food systems (5, 49, 50), further substantiating our motivation.

While our study was deliberately designed to control for seasonality and thus represents a worst-case, but most realistic, scenario in terms of what would be useful for application in anomaly detection in the industry, our results indicate that under such circumstances, explainable AI applied to microbiome data might become a valuable tool for monitoring and anomaly detection in food systems in the future.

### Raw milk microbial composition is dominated by few typical milk genera and overall uniform across sample types

To the best of our knowledge, this study characterized raw milk metagenomes in more sequencing depth than any other published work to date and demonstrates that there is a set of consensus microbes that were found to be stable elements across samples. We observed 32 microbial genera as present (RPM > 0.1) in all samples (excluding low-diversity outliers), see Supplemen-tal Table S5. *Pseudomonas, Serratia, Cutibacterium*, and *Staphylococcus* were the most abundant on average. This is in agreement with most of the literature, with *Pseudomonas* and *Staphylococcus* being reported in many studies that characterized the raw milk microbiome (51, 21, 52, 53). Mastitis causing bacteria, including *Streptococcus sp*., *Staphylococcus sp*., and coliforms such as *E. coli* and *Klebsiella* have been reported in milk microbiome studies (54, 55, 56), — all of which were detected in our dataset. Consequently, our data along with previously published data indicate that bacteria such as *Pseudomonas*, and *Staphylococcus*, among others, represent the core bulk tank milk microbiome.

In our study *Cutibacterium* was one of the most prevalent genera. *Cutibacterium* belongs to the family *Propionibacteriaceae*, which have been reported in the teat apex microbiota (51). A few skin-associated *Propionibacteria* have been reclassified as *Cutibacterium* (57), indicating that the teat skin microbiota might contribute to the bulk tank milk microbiota.

### Traditional microbial diversity metrics, clustering, and MDS and methods fail to differentiate sample classes

For the detection of anomalous samples, we evaluated several traditional methods for comparative analysis of the microbiome community. In this study clustering, cPCA, and MDS analyses did not indicate a strong separation of the three different classes of samples (baseline, outside farm, and antibiotics). This is in contrast with recently published studies (21, 22), in which milk samples were significantly different based on origin as identified by PERMANOVA and visualized with Principal Component Analysis. One explanation for the disparity is that in those studies, sampling was performed across seasons, thus increasing the variability between sample sites. In this study, we purposely sampled within a short time frame to control for seasonal variability, and thus avoid that confounder in our analyses. While our PERMANOVA results indicate that there was a significant difference in beta diversity between the three classes with multidimensional scaling (*p* = 0.0064), no clear separation could be observed between sample classes when the first two MDS are plotted. One might argue that most antibiotic samples are closer to one another than the other two classes, and thus that might explain our significant results within PERMANOVA. The high degree of uniformity observed here might be explained by our sampling strategy which was to constrain sampling to a short period of time to control for seasonality. It is also possible that management practices may explain differences between baseline and outside farm samples in our study.

### Bacterial taxa can be used as biomarkers for anomaly detection

Despite the overall uniformity in our data, three genera were identified to be differentially abundant between baseline and anomalous samples. *Enterobacter* have been reported to be present in milk microbiomes in many studies (51, 21, 52, 53), and *Coxiella* and *Morganella* were identified in this study but not in many others, perhaps because of to the large sequencing depth applied here. Nevertheless, these are environmental organisms that could be found in any farm, and their relative abundances could be informative when attempting to identify anomalies in milk.

In the case of milk received from an outside farm, we aimed to identify if the microbiome could signal more subtle differences since major influences, e.g. seasonality, region, and temperature, were all constant. Here while the cows and bulk tanks may be similar across farms, there were differences in farm management, diet, and milking protocols that one would expect to impart differences in the bulk tank milk microbiome. While baseline samples and outside farm samples were similar in their diversity and overall community membership, *Coxiella* was observed to have a significant increase in the outside farm samples. *Coxiella* is a known foodborne pathogen and a target organism for defining time and temperature combinations for milk pasteurization (58, 59). Thus it is not an unusual finding in raw milk microbiome, reiterating the fact that when using microbiome data for anomaly detection, one should not rely only on finding which features are unique to a sample class, but also how the relative abundances of features observed across classes might be associated with a particular class.

In milk samples spiked with milk from cows treated with antibiotics to control mastitis, the overall diversity does not differ from baseline diversity in a significant manner. Previous studies investigating bovine mastitis and antibiotic treatment have indicated that alpha diversity is significantly lower in mastitic milk when compared to healthy milk (60, 61). The lack of difference in diversity in our antibiotic anomalous samples is in accordance with the design of this study, in which antibiotic anomalies were not milk from an individual cow, but bulk tank milk spiked with 10% v/v of milk from an animal whose milk should not be entering the food supply chain. This represents a trace amount of contamination that although more difficult to detect, would represent a more realistic scenario. Even in such circumstances, we identified genera e.g. *Enterobacter* and *Morganella* that are observed with increased abundance in the antibiotic sample class. *Enterobacter spp*. are environmental mastitis-causing pathogens (54), hence expected in the mastitic milk from cows treated with antibiotics. *Morganella* is an environmental organism and part of the intestinal tract of mammals, and has been isolated from cheese(62). It can be an opportunistic pathogen and has been reported to infect calves (63).

### Explainable AI outperforms traditional methods in microbiome-based anomaly detection and can predict baseline vs. anomalous sample class using combined signal from all genera

For the detection of anomalous samples, we evaluated several traditional methods for comparative analysis of the microbiome community. Although clustering, cPCA, and MDS analyses did not indicate a strong separation of the three different classes of samples, predicting each type of anomaly versus the baseline could be done fairly accurately with both anomaly types using explainable AI even in our relatively small dataset. We observed that XGBoost was able to differentiate anomalous samples from baseline and were able to quantify the impact of the differentiating features with SHAP. Likewise, XGBoost, a gradient-boosted decision tree ensemble method, has also performed well in recent comparative studies on microbiome data (40, 29, 41).

While most machine learning algorithms function in a black-box manner, the explainability algorithm we used in this work— SHAP— assigns a value to each sample for each feature that describes the impact of that feature for predicting a specific sample class. One could speculate that this capability will be very useful for the food industry as evidence is amounting indicating that certain pathogens tend to co-occur with certain environmental microbes (64, 65). Thus, having the ability to “flag” microbes that predict a specific problematic sample class and might prove useful to inform sanitation and foodborne pathogen control practices. For example, *Listeria* has been shown to co-occur with certain taxa and thus observing those taxa as impactful genera in predicting a specific location might raise awareness about potential future problems with *Listeria* (64, 65).

The three genera observed to be most impactful by explainable AI were the same as those identified with the KS-test after Bonferroni correction; however by leveraging explainable AI, we were able to use the combined signal from all genera to assess the impact of the differing abundance of these microbes.

### Explainable AI can predict several fluid milk sample classes using an alternative datatype

Despite the high metagenomics sequencing depth, we recognize the small nature of the sample size (58 samples) when it pertains to testing machine learning algorithms (66, 67, 68), and that it may impart limitations on the quantitative and comparative assessments in characterizing the microbiomes of the multiple sample classes. We thus further validated our analysis approaches with publicly available bovine bulk tank milk data. The few publicly available bulk tank milk shotgun sequencing datasets had too low coverage (22, 69) to be appropriate for the application of our methodology. Thus, we performed our validation analysis based on publicly available fluid milk 16S amplicon sequencing (21, 70).

The results from our validation demonstrated a strong correlation with biological factors. Here, we observed that higher abundances of *Bacillus* and *Thermus*, and lower abundances of *Pseudomonas* and *Acinetobacter* influenced the prediction of post-pasteurization (Figure 7). These findings are consistent with the characteristics of the organisms that were found to have the most substantial impact on predicting post-pasteurization. Specifically, greater abundances of thermotolerant organisms such as *Bacillus* and *Thermus* were impactful for predicting post-pasteurization, as well as reduced abundances of typically thermosensitive *Pseudomonas*, were associated with prediction of post-pasteurization. Notably, these findings align with other analyses performed in the source study, including qPCR-based determination of cell numbers (23).

The milk microbiota has been reported to vary with season and stage of processing in several studies. Our validation analysis was also able to successfully predict milk samples by key attributes such as the season or the transport stage a milk sample was collected with high accuracy. Taken together, our results provide evidence for the feasibility of this approach and indicate that explainable AI has the potential to become a useful tool for microbiome-based quality monitoring for the food industry.

### Conclusion and future directions

In raw milk and other food systems, microbes can present important challenges to food safety in the case of pathogenic organisms and affect food quality such as flavor and storage attributes. Characterizing the microbial composition in a diverse set of food ingredients and products is of the utmost importance to better understand and improve the safety and quality of food. Since the microbiome is sensitive to changes in temperature, salinity, pH, and the composition of the material that it resides on among other things, it can also be utilized as an indicator for when food items deviate from a baseline of normality. For this study, our goal was to infer insights about each anomaly type compared to the baseline. As the number of samples collected was limited from a machine learning perspective, our intent was not to build a general machine learning model. Our intent was, instead, to investigate the potential use of an interpretable machine learning method (e.g., SHAP) to infer associations between microbial abundance and different sample types (baseline vs outside farm, baseline vs antibiotic-treated) and compare the ability of interpretable machine learning and traditional standard microbiome community analyses techniques to identify different sample types (and hence different sources of raw materials that could be used in food production). We therefore focus on the explanations provided by SHAP rather than the accuracy, the stability, or the generalizability of our machine learning model. Our overall aim is to provide a ‘proof-of-concept’ for this type of data and application. However, for results to be applicable for industry the sampling needs to be larger, therefore we envision to extend the approach to larger datasets as they become available. We demonstrate here that application of explainable AI applied to microbiome sequencing data could become a useful approach for anomaly detection in the food industry, particularly as sequencing technologies become more cost effective, laboratory processes are streamlined, and larger datasets are produced. Future challenges will include the need to define the appropriate specificity and sensitivity for models that can identify abnormalities in products or ingredients. Importantly, acceptable specificity and sensitivity will differ by the actual supply chains, and by factors such as the cost of false positive and false negatives, including the ease and cost of follow-up actions, e.g. follow up facility inspections or tests to determine whether detection of a microbiome abnormality represents an actual fraud or food safety incident.

## MATERIALS AND METHODS

### Milk sampling

Baseline raw bulk tank milk samples were collected daily from the Cornell University Ruminant Center (CURC; Hartford, NY) between September 5th and October 7th, 2018 and are referred to by the abbreviation BL. Anomalous samples from an outside farm (abbreviated as OF) were collected from a collaborating commercial dairy farm located in the same region (< 30 miles) and over the same time period as the Cornell Dairy. Bulk tank milk samples were collected aseptically into sterile 10 oz. vials (Capitol Plastics, Amsterdam, NY) and transported on ice to the Milk Quality Improvement Laboratory (Ithaca, NY). Anomalous samples from antibiotic-treated cows (abbreviated as ABX) were prepared at the laboratory by spiking the baseline sample of that day with 10% v/v with milk from an animal currently being treated with antibiotics, which was collected through a milking system collection device into a 10 oz vial.

Milk samples were aliquoted at the laboratory and frozen at -80°C until DNA extraction. A volume of 200 µL of milk samples were used as starting material for magnetic-based DNA extraction using a 96-well plate and CORE kit in a KingFisher instrument (Thermo Fisher Scientific, San Jose, CA, United States). Negative DNA extraction controls (reagents only) were carried out within the same plate for quality control. Extracted DNA was frozen at -80°C until library preparation and sequencing.

### Shotgun metagenome sequencing

Samples were quantified with a Qubit (Thermo Fisher Scientific, San Jose, CA, United States) before library preparation. Ten nanograms of each Qubit quantified genomic DNA was sheared with a Covaris E220 instrument operating SonoLab v6.2.6 generating approximately 300 bp DNA fragments according to the manufacturer’s protocol. Between 10 and 100 ng of fragmented DNA was processed into Illumina compatible sequencing libraries using sparQ DNA Library Prep Kit (Quantabio, Beverly, MA, United States). Each library was barcoded with unique dual index sequences (NEXTFLEX® Unique Dual Index Barcodes, BioO Scientific). Library size and amount were confirmed with a Bioanalyzer High Sensitivity DNA chip. Polymerase chain reaction primers and reagents included in the sparQ kit were used to perform PCR, and products were purified with AMPure XP beads. Equimolar libraries were pooled and subjected to Illumina NovaSeq 6000 sequencing at 2 × 150 bp (Illumina, San Diego, CA, United States). Shotgun whole metagenome sequencing was performed at the Genome Sciences and Bioinformatics Core at the Pennsylvania State University College of Medicine, Hershey, PA, United States. Illumina bcl2fastq (released version 2.20.0.422) was used to extract de-multiplexed sequencing reads.

### Read quality control and host filtering

Reads that included full length auxiliary sequences (junction adapter) P5 or P7 (“CTGTCTCTTATACACATCTCCGAGCCCAC-GAGAC” or “CTGTCTCTTATACACATCTGACGCTGCCGACGA”) were removed with a custom script (as were their read pairs), since their presence indicates issues with sequencing those particular reads (see Figure 2(b-d) in Illumina’s Sequencing Technical Note (71)). Trailing stretches of “G”, prevalent in Nextera sequences due to the two color channel technology, were removed with a custom script. If the length of the “G” tail was 30 or more, the read and its pair were discarded as low quality. The sequenced reads were then processed with TrimGalore (31) for adapter and quality trimming (parameters: –trim-n –paired –length 50 –phred33). The reads were handled as pairs through all the quality control and filtering steps.

Read filtering against the internal control PhiX and common host and contaminant genomes was performed with Kraken (32) as in Beck et al (5). PhiX reads were filtered against the NCBI Reference Sequence: NC_001422.1. Host reads were filtered against a database of plant and animal sequences introduced previously for metagenomic studies of food (5), which has been open-sourced and is available via PrecisionFDA https://precision.fda.gov/home/assets/file-GFfjPQj0ZqJV93P00bJ3vFgG-1. Additionally, kraken-filter with score threshold 0.1 was applied to avoid removing microbial reads.

### Microbial genus profiling

The reads passing quality control were classified as in Beck et al (5), against the NCBI’s RefSeq Complete (38) genome collection and corresponding taxonomy of bacterial, archaeal, viral, and eukaryotic microorganisms (approx. 7,800 genomes retrieved April 2017). Kraken (32) was used with a minimum score threshold of 0.05. Classified read counts per genus were collected as the sum of the read counts assigned to a genus or a taxonomic level below it. Sequencing blanks were used as negative controls to remove contaminating genera with the decontam R package (37) with the following parameters: threshold = 0.5 and normalize = TRUE. From this analysis, there were 14 genera which were removed from subsequent analysis: *Histophilus, Rahnella, Raoultella, T4virus, Pragia, C2virus, Methylophilus, Oceanobacillus, Streptosporangium, Fluviicola, Oenococcus, Alkalilimnicola, Geminocystis*, and *Brevibacillus*. Finally, classified reads per million quality-controlled sequenced reads (RPM) were computed for each genus and a threshold of 0.1 RPM applied to define *supported* genera, as described in Beck et al. (5). While the sequenced read depth was sufficient for genus-level taxonomic classification, it did not permit thorough gene or functional analysis.

### Community diversity

Shannon diversity was calculated from the supported microbial genera table using the diversity function in the vegan R package (72) with default parameters. Beta diversity was calculated using principles of compositional data analysis (73, 74). Therefore, read counts assigned to each genus were pseudo-counted by adding one in advance of computation of RPM prior to calculating the Aitchison distance from the microbial table. Beta diversity was calculated using the R package robCompositions (75) and hierarchical clustering was performed using base R function hclust using the “ward.D2” method.

Contrastive PCA (39) Python implementation was run with the aim of identifying enriched patterns in outside farm and antibiotic treated samples by contrasting them with baseline samples, on the supported microbial genera (Supplemental Table S4). In our target (foreground) data we kept OF, ABX and BL samples. In our background dataset, we only kept the baseline samples. We removed the 6 low diversity outliers from both target and background dataset. This was in an effort to uncover components which have high variance in the target dataset but low variance in the background dataset. We tried automatic assignment of alpha values where the algorithm generates and evaluates different alpha values and we also experimented with increasing alpha values systematically.

Multidimensional scaling (MDS, Matlab function cmdscale, p=2) and permutational multivariate analysis of variance (PERMANOVA, function f_permanova, iter=10,000, from the Fathom toolbox (76) for MATLAB) were applied on the pairwise Aitchison distances of all samples excluding the four baseline and two outside farm samples identified as low-diversity outliers, on the supported microbial genera table.

### Differences between baseline and outside farm, antibiotic treated samples

Two-sample statistical tests of individual features and corresponding visualizations on labeled data can be valuable as additional information to further support the results from explainable AI analysis, as done here. However, using two sample statistical tests alone only identifies significant differences for one single taxon, between two samples at a time and does not allow for assignment of a new sample to a class, as the RPM distributions per class are overlapping. Two-sample Kolmogorov-Smirnov tests (MATLAB function kstest2) were performed for each genus to determine microbes with significant differential abundance (p < 0.001) between baseline and each type of anomaly. Bonferroni multiple-comparison correction was applied, *p*^′^ = *pm*, where *m* is the number of genera, to obtain adjusted p-values *p*^′^. The differentially abundant genera were visualized using Violin Plot (77) in MATLAB.

### Explainable AI

To perform our explainable AI analysis we utilized the open source software ‘AutoXAI4Omics’, an automated explainable end-to-end ML tool developed for ‘omics datatypes (https://github.com/IBM/AutoXAI4Omics) (29).

For all datasets and classification tasks, we used AutoXAI4Omics to train and tune a series of ML models (XGBoost, Random Forest (RF), Support Vector Machines, Adaboost, K-Nearest Neighbors (KNN), LightGBM, Decision Trees, Extra Trees, Gradient Boosting, Stochastic Gradient Descent) using a train-test split ratio of 80:20. Hypertuning was performed on the training data using five-fold cross validation. For each classification task, predictive performance of all hyper-tuned models was assessed automatically by AutoXAI4Omics using the F1-score metric, and the top performing model was selected. Labels for Season and Processing Stage experiments were used that met the quality control and filtering criteria, with the exception of the Transport experiment. For the Transport experiment, sub-sampling was employed to randomly select samples, ensuring the class labels were more evenly balanced.

We used AutoXAI4Omics to apply an explainable AI algorithm called SHapley Additive exPlanations (SHAP), due to its ability to work with any machine learning model: tree-based models, such as XGBoost and LightGBM, as well as kernel-based and deep learning models. The explainability algorithm, SHAP, provides local explanations, i.e., interpretations of how the model predicts a particular value for a given sample. The local explanations show how each feature is contributing, either positively or negatively, to the prediction of a particular instance, for example of a particular class in case of classification task. After each models performance was evaluated, as described above, the top performing model cross-validation results were interpreted using SHAP to identify features which contribute most to the prediction.

We used the tuned top performing model coupled up with SHAP to explain the predictions (e.g., baseline vs anomalous) for each sample across the entire dataset. In addition to providing the ranked list of important features for a ML model, an advantage of SHAP over other feature importance methods is that it also explains how each of these impactful features is contributing (positively or negatively) to the prediction of specific phenotypic values. If we consider a binary classification task, the SHAP explainer returns two Shapley values tables of the same dimension of the input table (number of samples x number of genera/features), respectively for the class 0 (baseline) and the class 1 (anomalous). If we examine the table for class-0 baseline, each entry in the table is the SHAP impact (positive or negative) that a given genus has for the prediction of class baseline for a given sample. The absolute SHAP impact values for each feature are then averaged out across the entire set of samples to get an in-dication of the overall impact of a feature for the prediction, which results in a *ranked list of the most impactful features*.

We used AutoXAI4Omics to produce several plots providing visualisations of local explanations as well as a global view of local explanations that allow for interpretation of the entire model. The *SHAP beeswarm plot*, in particular, is a visualisation of the Shapley values matrix for a particular class (e.g., baseline). The plot shows the impact that each feature has on the prediction of the class for samples that share similar feature values. The y-axis is the ordered list (descending order of importance) of impactful features in predicting the class using a specific ML model (e.g., XGBoost). Therefore each row is a feature. The dots in each row are the data points, or samples, and are colored by the original feature value, that in this case is the genus abundance. The x-axis is the SHAP value or impact. A positive SHAP value/impact of a feature for a sample (the dot is on the right side of the x-axis) indicates that the feature (e.g., genus) has a positive impact in predicting the class (e.g., baseline), while a negative SHAP impact (the dot is on the left side of the x-axis) indicated that the feature has a negative impact on the prediction of the class. For each row (feature) the yellow and green dots can form separate clusters that are positioned towards the right or left side of the x-axis. This indicates that overall the feature (e.g., genus) tend to have a similar impact (positive or negative) for samples in which it has similar feature values (e.g., high or low abundance).

### Validation of anomaly detection in amplicon metagenomic samples

We further validated our findings by applying our explainable AI approach to publicly available datasets relevant to the dairy industry. Specifically, we selected two publicly available datasets from studies investigating the microbial profile of fluid milk using 16S rRNA amplicon sequencing for three comparisons. The data were retrieved from the European Nucleotide Archive (ERP015209, ERP114733) and contained 1,507 and 626 16S rRNA samples respectively (20, 23). Data were retrieved as .fasta files and subjected to a uniform pipeline using the DADA2 algorithm (78) in R and taxonomy was assigned using the SILVA (79) database. A count table was generated and was used to investigate the suitability of our explainable AI approach to distinguish between sample classes using amplicon-based data.

## Data availability

The sequencing data generated in this study are available at the NCBI BioProject PRJNA726965 https://www.ncbi.nlm.nih.gov/bioproject/PRJNA726965.

The code used to generate analyses in this study is available at https://github.com/gandalab/milk-anomaly-detection.

## CRediT author statement

Conceptualization (MW, KLB, NH, BK, EG), Methodology (EG, MW, KLB, NH), Resources (MW, NH, KLB, BK, MM, JK), Formal analysis (KLB, NH, BK, AA, APC), Data curation (VP), Validation (MM, JK), Visualization (KLB, NH, BK, AA, APC, MM, JK), Writing - Original Draft (KLB, NH), Writing - Review and Editing (KLB, NH, EG, MW).

## SUPPLEMENTAL MATERIAL

**FIG S1**. Sampling scheme.

**FIG S2**. Read counts per sample.

**FIG S3**. Contrastive PCA results.

**FIG S4**. Explainable AI results.

**TABLE S1**. Metadata on the raw milk samples.

**TABLE S2**. Summary of read counts.

**TABLE S3**. Read counts per genus.

**TABLE S4**. Supported genera RPM with contaminants removed.

**TABLE S5**. Core genera RPM with contaminants and outliers removed.

## ACKNOWLEDGMENTS

The authors would like to thank David Chambliss at IBM Research for his support with read filtering and data processing. This work was supported by the USDA National Institute of Food and Agriculture and Hatch Appropriations under Project #PEN04752, Accession #1023328 awarded to Erika Ganda. Sequencing was partially supported by a gift from IBM to Cornell University.

## REFERENCES

1. Spink J, Moyer DC. 11 2011. Defining the public health threat of food fraud. J food science 76. doi:10.1111/J.1750-3841.2011.02417.X.

2. Barnett J, Begen F, Howes S, Regan A, McConnon A, Marcu A, Rowntree S, Verbeke W. 1 2016. Consumers’ confidence reflections and response strategies following the horsemeat incident. Food Control 59:721–730. doi:10.1016/J.FOODCONT.2015.06.021.

3. Brooks S, Elliott CT, Spence M, Walsh C, Dean M. 2017. Four years post-horsegate: an update of measures and actions put in place following the horsemeat incident of 2013. NPJ science food 1. doi: 10.1038/S41538-017-0007-Z.

4. Handford CE, Campbell K, Elliott CT. 2016. Impacts of Milk Fraud on Food Safety and Nutrition with Special Emphasis on Developing Countries. Compr Rev Food Sci Food Saf 15 (1):130–142. doi: 10.1111/1541-4337.12181.

5. Beck KL, Haiminen N, Chambliss D, Edlund S, Kunitomi M, Huang BC, Kong N, Ganesan B, Baker R, Markwell P, Kawas B, Davis M, Prill RJ, Krishnareddy H, Seabolt E, Marlowe CH, Pierre S, Quintanar A, Parida L, Dubois G, Kaufman J, Weimer BC. 12 2021. Monitoring the microbiome for food safety and quality using deep shotgun sequencing. npj Sci Food 5 (1). doi:10.1038/s41538-020-00083-y.

6. Haiminen N, Edlund S, Chambliss D, Kunitomi M, Weimer BC, Ganesan B, Baker R, Markwell P, Davis M, Huang BC, Kong N, Prill RJ, Marlowe CH, Quintanar A, Pierre S, Dubois G, Kaufman JH, Parida L, Beck KL. 2019. Food authentication from shotgun sequencing reads with an application on high protein powders. npj Sci Food 3 (24). doi:10.1038/s41538-019-0056-6.

7. Martins G, Miot-Sertier C, Lauga B, Claisse O, Lonvaud-Funel A, Soulas G, Masneuf-Pomarède I. 8 2012. Grape berry bacterial microbiota: Impact of the ripening process and the farming system. Int J Food Microbiol 158:93–100. doi: 10.1016/J.IJFOODMICRO.2012.06.013.

8. Droby S, Wisniewski M. 6 2018. The fruit microbiome: A new frontier for postharvest biocontrol and postharvest biology. Postharvest Biol Technol 140:107–112. doi:10.1016/J.POSTHARVBIO.2018.03.004.

9. Walsh AM, Crispie F, Kilcawley K, O’Sullivan O, O’Sullivan MG, Claesson MJ, Cotter PD. 10 2016. Microbial Succession and Flavor Production in the Fermented Dairy Beverage Kefir. mSystems 1. doi: 10.1128/mSystems.00052-16.

10. Walsh AM, Crispie F, O’Sullivan O, Finnegan L, Claesson MJ, Cotter PD. 12 2018. Species classifier choice is a key consideration when analysing low-complexity food microbiome data. Microbiome 6:50. doi:10.1186/s40168-018-0437-0.

11. Duru IC, Laine P, Andreevskaya M, Paulin L, Kananen S, Tynkkynen S, Auvinen P, Smolander OP. 9 2018. Metagenomic and meta-transcriptomic analysis of the microbial community in Swiss-type Maasdam cheese during ripening. Int J Food Microbiol 281:10–22. doi:10.1016/J.IJFOODMICRO.2018.05.017.

12. Williams AG, Choi SC, Banks JM. 1 2002. Variability of the species and strain phenotype composition of the non-starter lactic acid bacterial population of cheddar cheese manufactured in a commercial creamery. Food Res Int 35:483–493. doi:10.1016/S0963-9969(01)00147-8.

13. Ranjan R, Rani A, Metwally A, McGee HS, Perkins DL. 2015. Analysis of the microbiome: Advantages of whole genome shotgun versus 16S amplicon sequencing. Biochem Biophys Res Commun doi: 10.1016/j.bbrc.2015.12.083.

14. Vangay P, Burgin J, Johnston A, Beck KL, Berrios DC, Blumberg K, Canon S, Chain P, Chandonia JM, Christianson D, Costes SV, Damerow J, Duncan WD, Dundore-Arias JP, Fagnan K, Galazka JM, Gibbons SM, Hays D, Hervey J, Hu B, Hurwitz BL, Jaiswal P, Joachimiak MP, Kinkel L, Ladau J, Martin SL, McCue LA, Miller K, Mouncey N, Mungall C, Pafilis E, Reddy TBK, Richardson L, Roux S, Schriml LM, Shaffer JP, Sundaramurthi JC, Thompson LR, Timme RE, Zheng J, Wood-Charlson EM, Eloe-Fadrosh EA. 2 2021. Microbiome Metadata Standards: Report of the National Microbiome Data Collaborative’s Workshop and Follow-On Activities. mSystems 6. doi: 10.1128/MSYSTEMS.01194-20.

15. Human Microbiome Project Consortium. 2012. A framework for human microbiome research. Nature 486:215–221. doi:10.1038/nature11209.

16. Greathouse KL, Sinha R, Vogtmann E. 2019. DNA extraction for human microbiome studies: The issue of standardization. Genome Biol 20:1–4. doi:10.1186/s13059-019-1843-8.

17. Costea PI, Zeller G, Sunagawa S, Pelletier E, Alberti A, Levenez F, Tramontano M, Driessen M, Hercog R, Jung FE, Kultima JR, Hayward MR, Coelho LP, Allen-Vercoe E, Bertrand L, Blaut M, Brown JRM, Carton T, Cools-Portier S, Daigneault M, Derrien M, Druesne A, Vos WMD, Finlay BB, Flint HJ, Guarner F, Hattori M, Heilig H, Luna RA, Vlieg JVH, Junick J, Klymiuk I, Langella P, Chatelier EL, Mai V, Manichanh C, Martin JC, Mery C, Morita H, O’toole PW, Orvain C, Patil KR, Penders J, Persson S, Pons N, Popova M, Salonen A, Saulnier D, Scott KP, Singh B, Slezak K, Veiga P, Versalovic J, Zhao L, Zoetendal EG, Ehrlich SD, Dore J, Bork P. 10 2017. Towards standards for human fecal sample processing in metagenomic studies. Nat Biotechnol 2017 35:11 35:1069–1076. doi:10.1038/nbt.3960.

18. Thompson L, Sanders J, McDonald D, Amir A, Ladau J, Locey K, Prill R, Tripathi A, Gibbons S, Ackermann G, Navas-Molina J, Janssen S, Kopylova E, Vázquez-Baeza Y, González A, Morton J, Mirarab S, Zech Xu Z, Jiang L, Haroon M, Kanbar J, Zhu Q, Jin Song S, Kosciolek T, Bokulich N, Lefler J, Brislawn C, Humphrey G, Owens S, Hampton-Marcell J, Berg-Lyons D, McKenzie V, Fierer N, Fuhrman J, Clauset A, Stevens R, Shade A, Pollard K, Goodwin K, Jansson J, Gilbert J, Knight R. 11 2017. A communal catalogue reveals Earth’s multiscale microbial diversity. Nat 2017 551:7681 551:457–463. doi: 10.1038/nature24621.

19. Ganda E, Beck KL, Haiminen N, Silverman JD, Kawas B, Cronk BD, Anderson RR, Goodman LB, Wiedmann M, Gibbons SM. 2021. DNA Extraction and Host Depletion Methods Significantly Impact and Potentially Bias Bacterial Detection in a Biological Fluid. mSystems 6 (3):e00619–21. doi:10.1128/mSystems.00619-21.

20. Kable ME, Srisengfa Y, Laird M, Zaragoza J, McLeod J, Heidenreich J, Marco ML. 2016. The core and seasonal microbiota of raw bovine milk in tanker trucks and the impact of transfer to a milk processing facility. mBio 7:1–13. doi:10.1128/mBio.00836-16.

21. Skeie SB, Håland M, Thorsen IM, Narvhus J, Porcellato D. 2019. Bulk tank raw milk microbiota differs within and between farms: A moving goalpost challenging quality control. J Dairy Sci 102:1959– 1971. doi:10.3168/JDS.2017-14083.

22. Yap M, Gleeson D, O’Toole PW, O’Sullivan O, Cotter PD, Björkroth J. 2021. Seasonality and Geography Have a Greater Influence than the Use of Chlorine-Based Cleaning Agents on the Microbiota of Bulk Tank Raw Milk. Appl Environ Microbiol 87 (22):e01081–21. doi: 10.1128/AEM.01081-21.

23. Xue Z, Brooks JT, Quart Z, Stevens ET, Kable ME, Heidenreich J, McLeod J, Marco ML. 2021. Microbiota Assessments for the Identification and Confirmation of Slit Defect-Causing Bacteria in Milk and Cheddar Cheese. mSystems 6 (1):e01114–20. doi:10.1128/mSystems.01114-20.

24. Porcellato D, Aspholm M, Skeie SB, Monshaugen M, Brendehaug J, Mellegård H. 2018. Microbial diversity of consumption milk during processing and storage. Int J Food Microbiol 266:21–30. doi: 10.1016/j.ijfoodmicro.2017.11.004.

25. McHugh AJ, Feehily C, Fenelon MA, Gleeson D, Hill C, Cotter PD, Gilbert JA. 2020. Tracking the Dairy Microbiota from Farm Bulk Tank to Skimmed Milk Powder. mSystems 5 (2):e00226–20. doi: 10.1128/mSystems.00226-20.

26. Gou W, Ling CW, He Y, Jiang Z, Fu Y, Fengzhe X, Miao ZL, Ting-yu S, Jie-Sheng L, Zhu H, Zhou HW, Chen YM, Zheng JS. 05 2020. Interpretable Machine Learning Algorithm Reveals Novel Gut Microbiome Features in Predicting Type 2 Diabetes. Curr Dev Nutr 4 (Supplement_2):1559–1559. doi:10.1093/cdn/nzaa062_016.

27. Wong CW, Yost SE, Lee JS, Gillece JD, Folkerts M, Reining L, Highlander SK, Eftekhari Z, Mortimer J, Yuan Y. 2021. Analysis of Gut Microbiome Using Explainable Machine Learning Predicts Risk of Diarrhea Associated With Tyrosine Kinase Inhibitor Neratinib: A Pilot Study. Front Oncol 11. doi:10.3389/fonc.2021.604584.

28. Liu Y, Mé G, Havulinna AS, Salomaa V. 2022. Early prediction of incident liver disease using conventional risk factors and gutmicrobiome-augmented gradient boosting. Cell Metab 34:719– 730.e4. doi:10.1016/j.cmet.2022.03.002.

29. Carrieri AP, Haiminen N, Maudsley-Barton S, Gardiner L, Murphy B, Mayes A, Paterson S, Grimshaw S, Winn M, Shand C, Hadjidoukas P, Rowe W, Hawkins S, MacGuire-Flanagan A, Tazzioli J, Kenny J, Parida L, Hoptroff M, Pyzer-Knapp E. 2 2021. Explainable AI reveals changes in skin microbiome composition linked to phenotypic differences. Sci Rep. 11 (4565). doi:10.1038/s41598-021-83922-6.

30. Andrews S. 2010. FastQC: A Quality Control Tool for High Throughput Sequence Data http://www.bioinformatics.babraham.ac.uk/projects/fastqc.

31. Krueger F. 2012. Trim Galore https://github.com/FelixKrueger/TrimGalore.

32. Wood DE, Salzberg SL. 2014. Kraken: ultrafast metagenomic sequence classification using exact alignments. Genome Biol 15 (3):R46. doi:10.1186/gb-2014-15-3-r46.

33. Salter SJ, Cox MJ, Turek EM, Calus ST, Cookson WO, Moffatt MF, Turner P, Parkhill J, Loman NJ, Walker AW. 11 2014. Reagent and laboratory contamination can critically impact sequence-based microbiome analyses. BMC biology 12. doi:10.1186/S12915-014-0087-Z.

34. Weyrich LS, Farrer AG, Eisenhofer R, Arriola LA, Young J, Selway CA, Handsley-Davis M, Adler CJ, Breen J, Cooper A. 7 2019. Laboratory contamination over time during low-biomass sample analysis. Mol ecology resources 19:982–996. doi:10.1111/1755-0998.13011.

35. Eisenhofer R, Minich JJ, Marotz C, Cooper A, Knight R, Weyrich LS. 2 2019. Contamination in Low Microbial Biomass Microbiome Studies: Issues and Recommendations. Trends microbiology 27:105–117. doi: 10.1016/J.TIM.2018.11.003.

36. Hornung BVH, Zwittink RD, Kuijper EJ. 5 2019. Issues and current standards of controls in microbiome research. FEMS microbiology ecology 95. doi:10.1093/FEMSEC/FIZ045.

37. Davis NM, Proctor DM, Holmes SP, Relman DA, Callahan BJ. 2018. Simple statistical identification and removal of contaminant sequences in marker-gene and metagenomics data. Microbiome 6 (226). doi:10.1186/S40168-018-0605-2/FIGURES/6.

38. O’Leary NA, Wright MW, Brister JR, Ciufo S, Haddad D, McVeigh R, Rajput B, Robbertse B, Smith-White B, Ako-Adjei D, Astashyn A, Badretdin A, Bao Y, Blinkova O, Brover V, Chetvernin V, Choi J, Cox E, Ermolaeva O, Farrell CM, Goldfarb T, Gupta T, Haft D, Hatcher E, Hlavina W, Joardar VS, Kodali VK, Li W, Maglott D, Masterson P, McGarvey KM, Murphy MR, O’Neill K, Pujar S, Rangwala SH, Rausch D, Riddick LD, Schoch C, Shkeda A, Storz SS, Sun H, Thibaud-Nissen F, Tolstoy I, Tully RE, Vatsan AR, Wallin C, Webb D, Wu W, Landrum MJ, Kimchi A, Tatusova T, DiCuccio M, Kitts P, Murphy TD, Pruitt KD. 11 2015. Reference sequence (RefSeq) database at NCBI: current status, taxonomic expansion, and functional annotation. Nucleic Acids Res 44 (D1):D733–D745. doi: 10.1093/nar/gkv1189.

39. Abid A, Zhang MJ, Bagaria VK, Zou J. 2018. Exploring patterns enriched in a dataset with contrastive principal component analysis. Nat communications 9 (1):2134.

40. Wang XW, Liu YY. 2020. Comparative study of classifiers for human microbiome data. Med Microecol 4:100013. doi: 10.1016/j.medmic.2020.100013.

41. Chen Y, Wang H, Lu W, Wu T, Yuan W, Zhu J, Lee YK, Zhao J, Zhang H, Chen W. 2022. Human gut microbiome aging clocks based on taxonomic and functional signatures through multi-view learning. Gut Microbes 14 (1):2025016. doi:10.1080/19490976.2021.2025016.

42. Lundberg SM, Lee SI. 2017. A unified approach to interpreting model predictions. Proc Neural Inf Process Syst 30:4765–4774.

43. Jiang L, Haiminen N, Carrieri AP, Huang S, Vázquez-Baeza Y, Parida L, Kim HC, Swafford AD, Knight R, Natarajan L. 2021. Utilizing stability criteria in choosing feature selection methods yields reproducible results in microbiome data. Biometrics doi: 10.1111/biom.13481.

44. Montgomery H, Haughey SA, Elliott CT. 9 2020. Recent food safety and fraud issues within the dairy supply chain (2015–2019). Glob Food Secur 26:100447. doi:10.1016/J.GFS.2020.100447.

45. FDA. 2017. Grade “A” Pasteurized Milk Ordinance.

46. Dini-Andreote F, Pylro VS, Baldrian P, Elsas JDV, Salles JF. 8 2016. Ecological succession reveals potential signatures of marine–terrestrial transition in salt marsh fungal communities. The ISME J 10:1984. doi:10.1038/ISMEJ.2015.254.

47. Leung H, Long X, Ni Y, Qian L, Nychas E, Siliceo SL, Pohl D, Hanhineva K, Liu Y, Xu A, Nielsen HB, Belda E, Clément K, Loomba R, Li H, Jia W, Panagiotou G. 6 2022. Risk assessment with gut microbiome and metabolite markers in NAFLD development. Sci Transl Med 14:855. doi:10.1126/scitranslmed.abk0855.

48. Dini-Andreote F, Elsas JDV, Olff H, Salles JF. 12 2018. Dispersal-competition tradeoff in microbiomes in the quest for land colonization. Sci Reports 8. doi:10.1038/S41598-018-27783-6.

49. Yap M, Ercolini D, Álvarez Ordóñez A, O’Toole PW, O’Sullivan O, Cotter PD. 2022. Next-Generation Food Research: Use of Meta-Omic Approaches for Characterizing Microbial Communities Along the Food Chain. Annu Rev Food Sci Technol 13 (1):361–384. doi: 10.1146/annurev-food-052720-010751.

50. Mukherjee A, Gómez-Sala B, O’Connor EM, Kenny JG, Cotter PD. 5 2022. Global Regulatory Frameworks for Fermented Foods: A Review. Anomaly detection in dairy with metagenome sequencing Front Nutr 0:1084. doi:10.3389/FNUT.2022.902642.

51. Derakhshani H, Fehr KB, Sepehri S, Francoz D, De Buck J, Barkema HW, Plaizier JC, Khafipour E. 2018. Invited review: Microbiota of the bovine udder: Contributing factors and potential implications for udder health and mastitis susceptibility. J Dairy Sci 101 (12):10605–10625. doi:10.3168/jds.2018-14860.

52. Liu J, Zhu Y, Jay-Russell M, Lemay DG, Mills DA. 2020. Reservoirs of antimicrobial resistance genes in retail raw milk. Microbiome 8 (1):99. doi:10.1186/s40168-020-00861-6.

53. Ruegg PL. 2022. The bovine milk microbiome – an evolving science. Domest Animal Endocrinol 79:106708. doi: 10.1016/j.domaniend.2021.106708.

54. Oliveira L, Hulland C, Ruegg PL. 2013. Characterization of clinical mastitis occurring in cows on 50 large dairy herds in Wisconsin. J Dairy Sci 96 (12):7538–7549. doi: 10.3168/jds.2012-6078.

55. Bhatt VD, Ahir VB, Koringa PG, Jakhesara SJ, Rank DN, Nauriyal DS, Kunjadia AP, Joshi CG. 2012. Milk microbiome signatures of subclinical mastitis-affected cattle analysed by shotgun sequencing. J Appl Microbiol 112 (4):639–650. doi: 10.1111/j.1365-2672.2012.05244.x.

56. Hoque MN, Istiaq A, Clement RA, Sultana M, Crandall KA, Siddiki AZ, Hossain MA. 9 2019. Metagenomic deep sequencing reveals association of microbiome signature with functional biases in bovine mastitis. Sci Reports 2019 9:1 9:1–14. doi:10.1038/s41598-019-49468-4.

57. Scholz CFP, Kilian M. 11 2016. The natural history of cutaneous propionibacteria, and reclassification of selected species within the genus propionibacterium to the proposed novel genera acidipropionibacterium gen. Nov., cutibacterium gen. nov. and pseudopropionibacterium gen. nov. Int J Syst Evol Microbiol 66:4422–4432. doi: 10.1099/IJSEM.0.001367/CITE/REFWORKS.

58. Commission CA, et al.. 2004. Code of hygienic practice for milk and milk products. CAC/RCP 57-2004. Food Agric Organ Rome.

59. Wittwer M, Hammer P, Runge M, Valentin-Weigand P, Neubauer H, Henning K, Mertens-Scholz K. 1 2022. Inactivation Kinetics of Coxiella burnetii During High-Temperature Short-Time Pasteurization of Milk. Front Microbiol 12:3975. doi:10.3389/fmicb.2021.753871.

60. Ganda EK, Bisinotto RS, Lima SF, Kronauer K, Decter DH, Oikonomou G, Schukken YH, Bicalho RC. 2016. Longitudinal metagenomic profiling of bovine milk to assess the impact of intramammary treatment using a third-generation cephalosporin. Sci Reports 6:1–13. doi:10.1038/srep37565.

61. Ganda EK, Gaeta N, Sipka A, Pomeroy B, Oikonomou G, Schukken YH, Bicalho RC. 2017. Normal milk microbiome is reestablished following experimental infection with Escherichia coli independent of intramammary antibiotic treatment with a thirdgeneration cephalosporin in bovines. Microbiome 5. doi: 10.1186/s40168-017-0291-5.

62. Ryser LT, Arias-Roth E, Perreten V, Irmler S, Bruggmann R. 2021. Genetic and Phenotypic Diversity of Morganella morganii Isolated From Cheese. Front Microbiol 12:3378. doi: 10.3389/fmicb.2021.738492.

63. Li G, Niu X, Yuan S, Liang L, Liu Y, Hu L, Liu J, Cheng Z. 12 2018. Emergence of Morganella morganii subsp. morganii in dairy calves, China. Emerg Microbes & Infect 7. doi:10.1038/S41426-018-0173-3.

64. Fox EM, Solomon K, Moore JE, Wall PG, Fanning S. 2014. Phylogenetic profiles of in-house microflora in drains at a food production facility: Comparison and biocontrol implications of listeria-positive and -negative bacterial populations. Appl Environ Microbiol 80:3369– 3374. doi:10.1128/AEM.00468-14.

65. Willis C, Jørgensen F, Aird H, Elviss N, Fox A, Jenkins C, Fenelon D, Sadler-Reeves L, McLauchlin J. 2 2018. An assessment of the microbiological quality and safety of raw drinking milk on retail sale in England. J Appl Microbiol 124:535–546. doi:10.1111/JAM.13660.

66. Cui Z, Gong G. 9 2018. The effect of machine learning regression algorithms and sample size on individualized behavioral prediction with functional connectivity features. NeuroImage 178:622–637. doi: 10.1016/j.neuroimage.2018.06.001.

67. Balki I, Amirabadi A, Levman J, Martel AL, Emersic Z, Meden B, Garcia-Pedrero A, Ramirez SC, Kong D, Moody AR, Tyrrell PN. 11 2019. Sample-Size Determination Methodologies for Machine Learning in Medical Imaging Research: A Systematic Review. Can Assoc Radiol journal = J l’Association canadienne des radiologistes 70:344– 353. doi:10.1016/J.CARJ.2019.06.002.

68. Vabalas A, Gowen E, Poliakoff E, Casson AJ. 11 2019. Machine learning algorithm validation with a limited sample size. PLoS ONE 14. doi: 10.1371/journal.pone.0224365.

69. Yap M, Feehily C, Walsh CJ, Fenelon M, Murphy EF, McAuliffe FM, van Sinderen D, O’Toole PW, O’Sullivan O, Cotter PD. Dec 2020. Evaluation of methods for the reduction of contaminating host reads when performing shotgun metagenomic sequencing of the milk microbiome. Sci Reports 10 (1):21665. doi: 10.1038/s41598-020-78773-6.

70. Sun L, Åse Lundh, Höjer A, Bernes G, Nilsson D, Johansson M, Hetta M, Gustafsson AH, Saedén KH, Dicksved J. 1 2022. Milking system and premilking routines have a strong effect on the microbial community in bulk tank milk. J dairy science 105:123–139. doi: 10.3168/JDS.2021-20661.

71. Illumina. 2012. Data Processing of Nextera® Mate Pair Reads on Illumina Sequencing Platforms https://www.illumina.com/documents/products/technotes/technote_nextera_mat

72. Oksanen J, Blanchet FG, Friendly M, Kindt R, Legendre P, McGlinn D, Minchin PR, O’Hara RB, Simpson GL, Solymos P, Stevens MHH, Szoecs E, Wagner H. 2020. vegan: Community Ecology Package. https://CRAN.R-project.org/package=vegan. R package version 2.5-7.

73. Gloor GB, Macklaim JM, Pawlowsky-Glahn V, Egozcue JJ. 11 2017. Microbiome Datasets Are Compositional: And This Is Not Optional. Front Microbiol 8:2224. doi:10.3389/fmicb.2017.02224.

74. Aitchison J, Barceló-Vidal C, Martín-Fernández JA, Pawlowsky-Glahn V. 2000. Logratio Analysis and Compositional Distance. Math Geol 32:271–275. doi:10.1023/A:1007529726302.

75. Templ M, Hron K, Filzmoser P. 7 2011. robCompositions: An R-package for Robust Statistical Analysis of Compositional Data. John Wiley & Sons, Ltd. doi:10.1002/9781119976462.ch25.

76. Jones DL. 2017. Fathom Toolbox for MATLAB: software for multivariate ecological and oceanographic data analysis https://www.usf.edu/marine-science/research/matlab-resources/index.aspx.

77. Hoffmann H. 2015. Violin Plot. MATLAB Central File Exchange https://www.mathworks.com/matlabcentral/fileexchange/45134-violin-plot.

78. Callahan BJ, McMurdie PJ, Rosen MJ, Han AW, Johnson AJ, Holmes SP. 2016. DADA2: High-resolution sample inference from Illumina amplicon data. Nat Methods 13:581–583. doi:10.1038/nmeth.3869. Callahan Benjamin JMcMurdie, Paul JRosen, Michael JHan, Andrew WJohnson, Amy Jo AHolmes, Susan PengR01 AI112401/AI/NIAID NIH HHS/2016/05/24 06:00 Nat Methods. 2016 Jul;13(7):581–3. doi: 10.1038/nmeth.3869. Epub 2016 May 23.

79. Quast C, Pruesse E, Yilmaz P, Gerken J, Schweer T, Yarza P, Peplies J, Glöckner FO. 1 2013. The SILVA ribosomal RNA gene database project: improved data processing and web-based tools. Nucleic acids research 41. doi:10.1093/NAR/GKS1219.

