## Supplemental Figures for "Development and evaluation of statistical and Artificial Intelligence approaches with microbial shotgun metagenomics data as an untargeted screening tool for use in food production"

### Detecting anomalous content in dairy with metagenomic sequencing

*Haiminen et al., 2022*

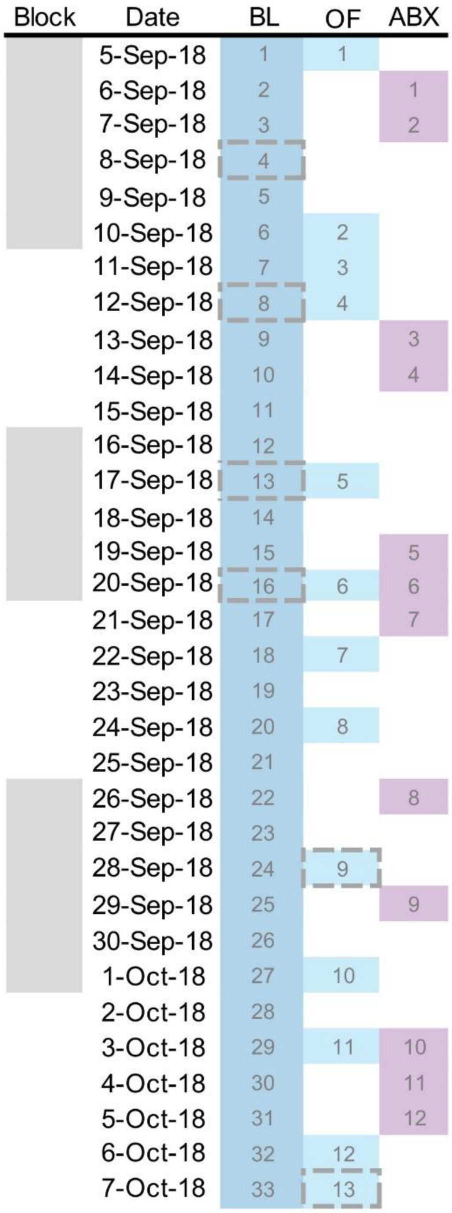


**Supplemental Figure S1:** Sampling Scheme. Dates were block randomized to ensure even distribution across the sampling period for each anomaly. Sampling occurred in a short time frame to control for potential season effect. Four baseline (BL) and two outside farm (OF) samples that were discarded as low-diversity outliers are indicated by grey dashed lines.

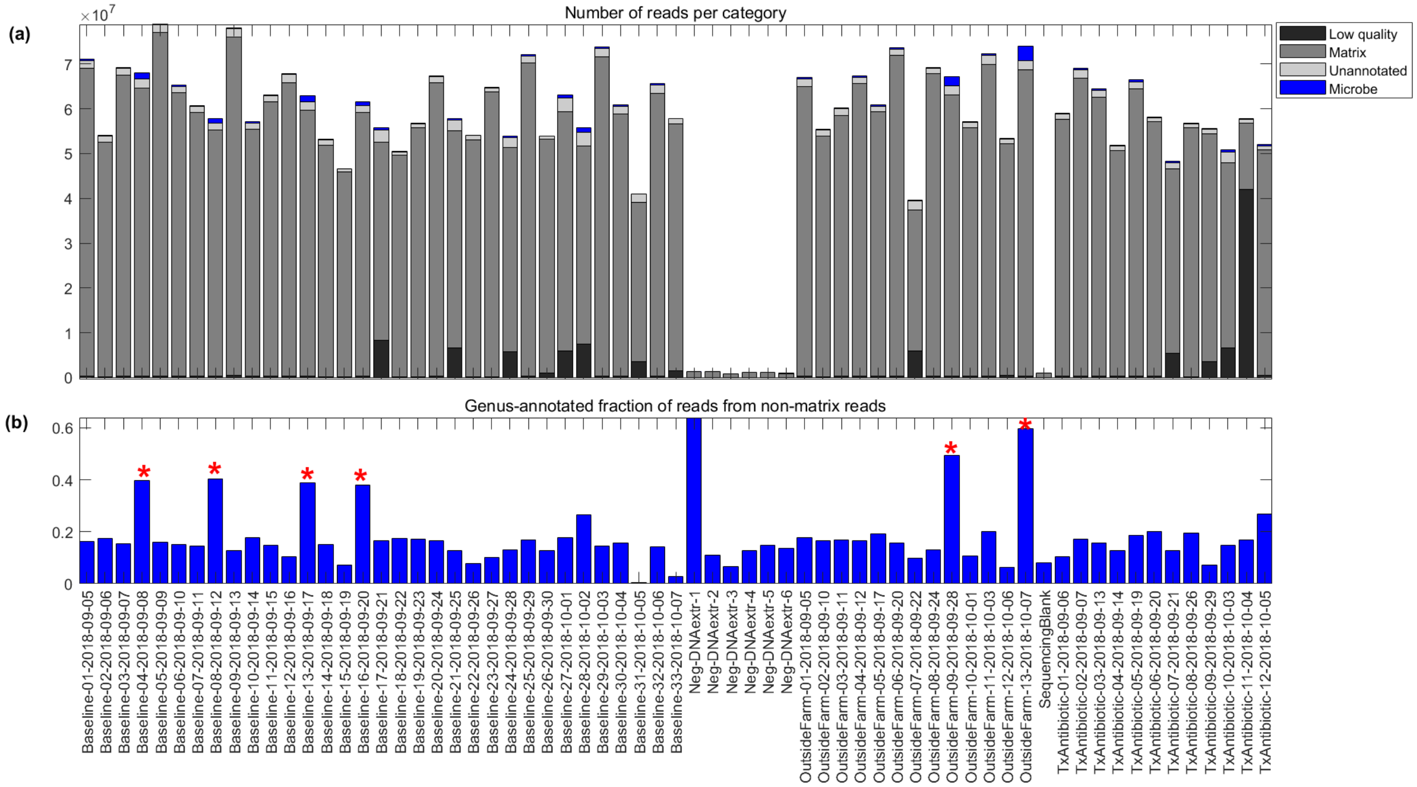


**Supplemental Figure S2:** Read counts per sample. **(a)** Total number of input reads per sample are shown, divided into low quality, matrix filtered, unannotated, and microbial genus-assigned (after removing contaminating genera). **(b)** The fraction of genus-annotated reads from the non-matrix reads with low-diversity outlier samples indicated with red stars.

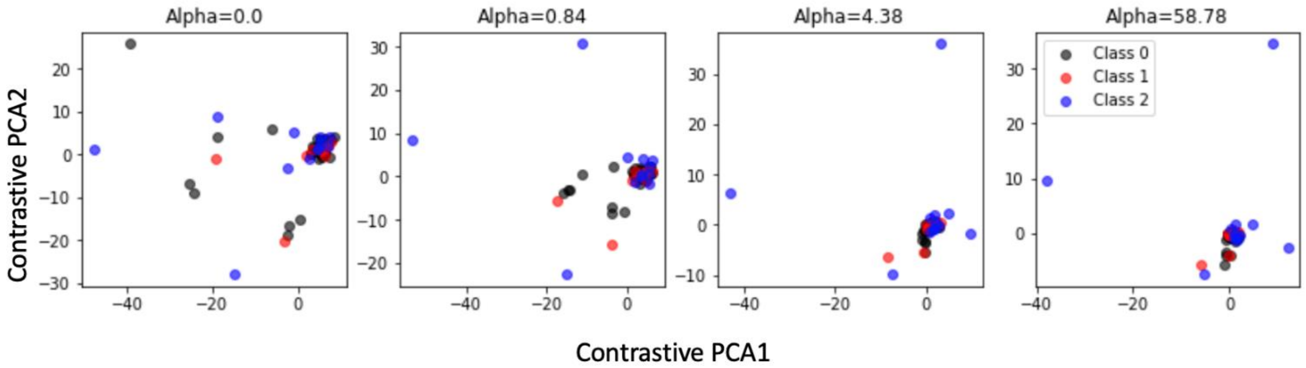


**Supplemental Figure S3:** Contrastive PCA results. Each box corresponds to a different value of alpha. Class 0 (black dots) indicate baseline samples (the background samples). Class 1 (red dots) indicate outside farm sample and class 2 (blue dots) indicate antibiotic treated samples.

### (a) Outside farm vs. baseline

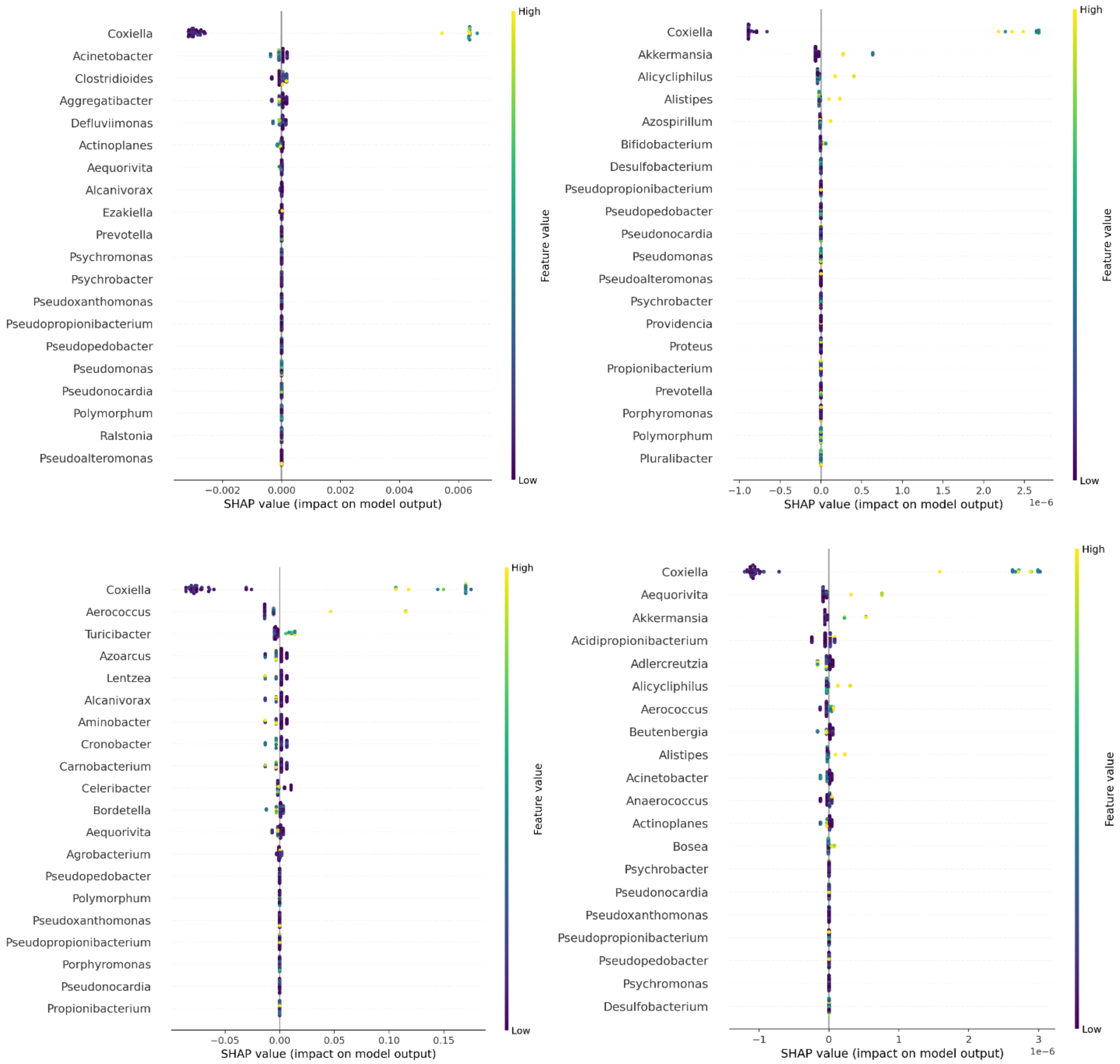

(b) Antibiotic treated vs. baseline

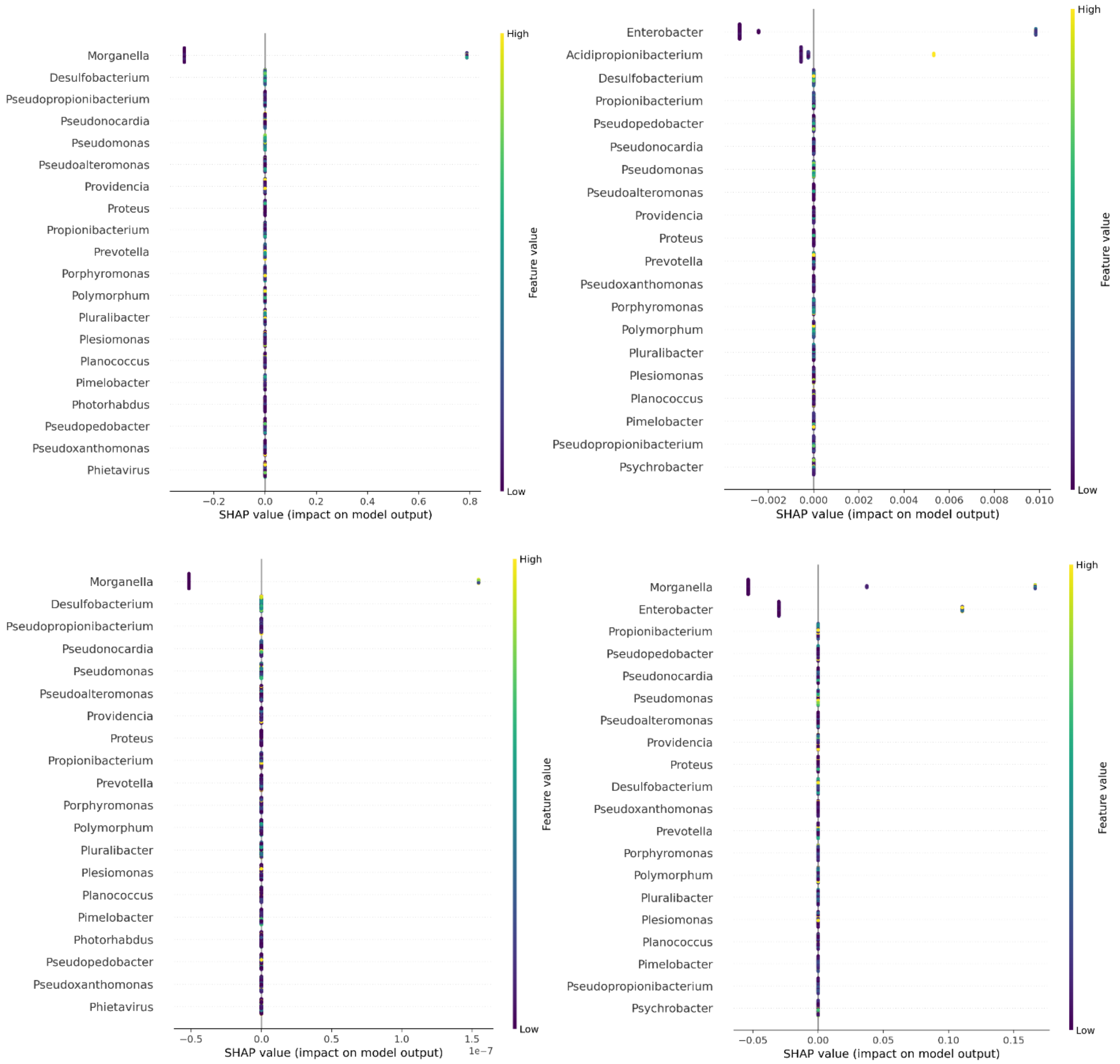


**Supplemental Figure S4**: SHAP results. SHAP dot plots for the most impactful features when predicting (a) Outside farm or (b) Antibiotic treated sample class vs. baseline in four independent iterations.
